# Bacterial genome engineering using CRISPR RNA-guided transposases

**DOI:** 10.1101/2023.03.18.533263

**Authors:** Diego R. Gelsinger, Phuc Leo H. Vo, Sanne E. Klompe, Carlotta Ronda, Harris Wang, Samuel H. Sternberg

## Abstract

CRISPR-associated transposons (CASTs) have the potential to transform the technology landscape for kilobase-scale genome engineering, by virtue of their ability to integrate large genetic payloads with high accuracy, easy programmability, and no requirement for homologous recombination machinery. These transposons encode efficient, CRISPR RNA-guided transposases that execute genomic insertions in *E. coli* at efficiencies approaching ∼100%, generate multiplexed edits when programmed with multiple guides, and function robustly in diverse Gram-negative bacterial species. Here we present a detailed protocol for engineering bacterial genomes using CAST systems, including guidelines on the available homologs and vectors, customization of guide RNAs and DNA payloads, selection of common delivery methods, and genotypic analysis of integration events. We further describe a computational crRNA design algorithm to avoid potential off-targets and CRISPR array cloning pipeline for DNA insertion multiplexing. Starting from available plasmid constructs, the isolation of clonal strains containing a novel genomic integration event-of-interest can be achieved in 1 week using standard molecular biology techniques.

## Introduction

The field of biology has been revolutionized by the discovery of adaptive immune systems encoded by clustered regularly interspaced short palindromic repeats (CRISPR) and CRISPR-associated (Cas) genes, and the subsequent harnessing of CRISPR-Cas systems for genome engineering. In particular, CRISPR-Cas based genetic manipulations have been applied to several model (i.e., *E. coli*) and non-model organisms across all domains of life with high efficiency^1–4^ which has greatly aided our understanding of the biology underlying these organisms. CRISPR-Cas systems encode a diverse repitoire of RNA-guided CRISPR-associated (Cas) effector nucleases that perform interference on invading mobile genetic elements. These programmable endonucleases (Cas) can target DNA or RNA for interference by complexing with a non-coding CRISPR RNA (crRNA, also referred to as a guide RNA), and have been repurposed as efficient genome editing tools that can be directed to nearly any DNA target sequence using a re-coded guide RNA^5–9^. The most prominent of these biotechnologies, CRISPR-Cas9, allows targeted cutting of double stranded DNA and has vastly expanded the eukaryotic genome engineering toolkit^10,11^. Despite their bacterial origins, conventional CRISPR-based approaches have not fundamentally changed the landscape of bacterial genome engineering to the same degree due to various limitations, such as cytotoxicity^12^.

Many bacterial engineering applications have instead utilized recombineering, a common method that relies on homologous recombination (HR) between genomic DNA and synthetic user-provided donor molecules containing the desired DNA insert flanked by homology arms^13,14^. In recent years, CRISPR-Cas9 has been combined with recombineering to provide counterselection against unedited cells via targeted cleavage of the wild-type allele, thereby enabling programmable scarless editing without the need for drug marker selection^15–19^. Recombineering strategies, while often effective, typically require the introduction of exogenous recombination machinery (e.g. Lambda red system), and can suffer from low efficiencies, particularly for the insertion of multi-kb DNA payloads^20^. Additionally, recombineering often translates poorly between diverse target species due to host specificity of exogenous recombination proteins^21^.

Recent advances in other areas of synthetic biology and genome engineering have provided fundamental avenues to further our understanding of bacterial cellular biology, pathogenicity, and functional genomics^22^. These advances have enabled the use of bacteria as diagnostic and therapeutic agents targeting diverse diseases^23^, and as “biofactories” for industrial production of biofuels and beneficial small molecules^24^. Many of these applications require genomic insertion of customized DNA payloads (i.e. “knock-ins”), which allow for stable maintenance of desired expression cassettes at predictable copy numbers and reduced metabolic burden, but without drug marker selection or population heterogeneity that are associated with plasmid-based expression^25^. However, existing transposase and recombinase platforms commonly applied for DNA insertion, such as Cre recombinase or Tn*7* transposase, recognize fixed target sequences and are thus not readily programmable^26–29^. Alternatively, integration systems such as the Tn*5* or Mariner transposase have little to no target specificity and catalyze insertion into random genomic sites^30,31^, which is undesirable for systematic and controllable strain engineering programs.

We recently described a powerful new addition to the genome engineering toolbox, in which we exploited CRISPR-associated transposons (CASTs) to achieve highly efficient, targeted DNA integration of large kilobase-scale payloads^32,33^ (**Fig. 1A**). These systems combine the ease of programmability of CRISPR-Cas systems, with the efficient chemistry afforded by transposase enzymes, to direct DSB-free and recombination-independent DNA insertion. Since their initial bioinformatic discovery^34^, we and others have harnessed CAST systems for a range of engineering applications in diverse microbial species^32,35–38^, taking advantage of their ability to integrate payloads with high efficiency, function independently of host factors, perform multiplexed insertions with multiple guide RNAs, and mobilize payloads ranging from less than 1 kb to more than 10 kb in size.

**Fig 1.**
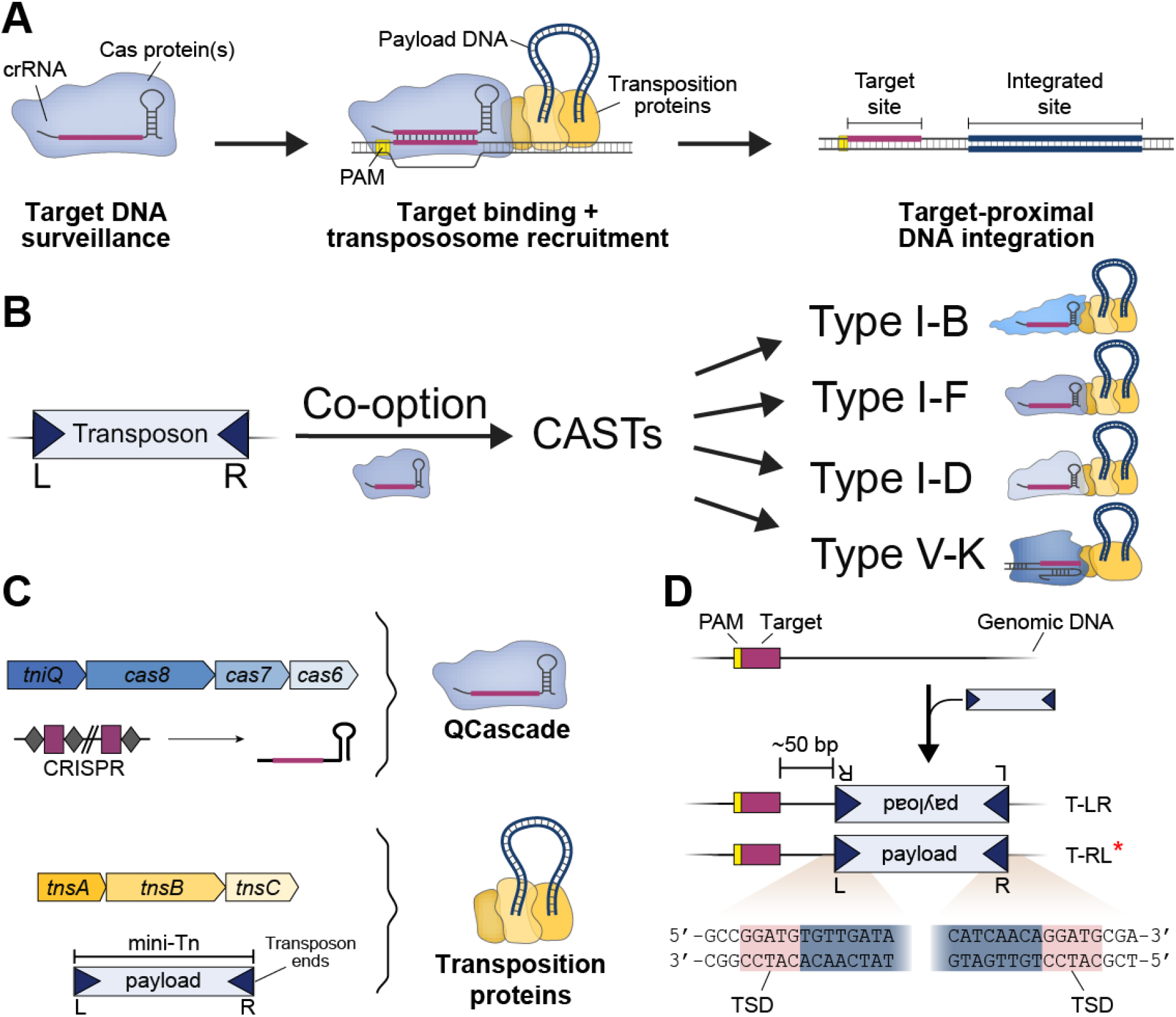
Overview of CRISPR-associated transposons (CASTs). (A) Simplified schematic of the general mechanism of RNA-guided DNA transposition. CRISPR-Cas effector complexes, consisting of a mature CRISPR RNA (crRNA) and one or more Cas proteins, recognize and bind genomic target sites using RNA-DNA complementarity. Subsequent recruitment of transposase proteins in complex with the donor DNA leads to integration of the mini-Tn a fixed distance downstream of the target site. The mini-Tn can be customized with user-defined payloads. (B) Tn7-like transposons have co-opted at least four different families of nuclease-deficient CRISPR-Cas systems during CAST evolution: Type I-B, I-D, I-F, and V-K. (C) Required components for RNA-guided DNA integration using Type I-F CASTs. The DNA binding complex, QCascade, consists of TniQ, multiple Cas proteins, and a mature crRNA that is processed by Cas6. TnsA and TnsB catalyze DNA excision and integration chemistry, aided by the mediator ATPase, TnsC. Mini-Tn substrates must be flanked by transposon right (R) and left (L) ends. (D) DNA insertions occur ∼50 bp downstream of the target site in one of two possible orientations, defined by which transposon end is closest to the target site, denoted T-LR and T-RL. T-RL products (*) are preferentially generated by Type I-F CASTs, and products exhibit hallmark 5-bp target-site duplications (TSD).

### Mechanism and development of Type I-F CASTs

CRISPR-associated transposons are evolutionarily diverse, having arisen from at least four independent exaptation events in which Tn7-like transposons repurposed nuclease-deficient CRISPR-Cas systems from Type I-B, Type I-D, Type I-F, or Type V-K (**Fig. 1B**)^33,39–44^. All CAST systems rely on the conserved DDE-family TnsB transposase to perform the concerted strand transfer reactions during transposition, alongside common accessory factors including TnsC and TniQ, but the molecular basis of DNA targeting differs: Type I CAST systems employ a multi-subunit RNA-guided DNA binding complex called Cascade for target selection, whereas Type V-K CAST systems employ the single-effector protein, Cas12k^33,39,40,42^. Importantly, other mechanistic parameters differ sharply between CAST systems, including the number of molecular components, purity of insertion products, genome-wide fidelity, and on-target efficiency^33,37,39,42,45,46^. Due to key advantages we previously reported for Type I-F CAST systems^32^, we prioritized subsequent technology development on these systems and focus the remainder of this protocol on providing technical details and guidelines for their use in bacteria. However, we point interested readers to recent studies that describe both mechanistic and technological advances in the study and application of Type V-K CAST systems^45,47,48^.

Natural CRISPR-associated transposons are bounded by conserved transposon left (L) and right (R) end sequences, and thus, genetic payloads intended for site-specific genomic insertion end by engineered CAST systems must also be encoded within a mini-transposon (mini-Tn) context bounded by the same sequence features. Two molecular machineries encoded within I-F CAST systems are essential for directing and catalyzing RNA-guided DNA transposition: an RNA-guided DNA targeting complex known as TniQ-Cascade (hereafter simply QCascade)— which comprises a CRISPR RNA (crRNA) guide and protein components TniQ, Cas8, Cas7, and Cas6^33,40^ — and the heteromeric transposase complex TnsABC, made up of the TnsA endonuclease, the TnsB transposase, and the TnsC ATPase^33,49^ (**Fig. 1C**). QCascade uses a 32-nt guide sequence to bind 32-bp DNA targets (*target site*) flanked by a 5’-CN-3’ PAM^33,40,41^, leading to integration of the mini-Tn at a fixed distance ∼50-bp downstream of the target site (*integration site*) defined by the molecular footprint of associated transposition proteins^33,46,50^ (**Fig. 1D**). Importantly, DNA integration does not disrupt the target site itself, leaving open the possibility that constitutive expression of CAST machinery could lead to iterative rounds of targeting and DNA insertion. However, these tandem insertions are rarely generated because of a feature intrinsic to Tn7-like transposons known as target immunity (discussed further below)^33,42,49,51–54^.

Type I-F CASTs generate simple insertion products through a non-replicative cut-and-paste reaction, whereas Type V-K CASTs lack the TnsA endonuclease and therefore generate cointegrate products through a replicative mechanism^46,55,56^. Transposition products feature hallmark target-site duplications (TSDs) flanking the inserted payload, in which 5-bp of genomic sequence is precisely duplicated (**Fig. 1D**). Orientation control is another key feature of RNA-guided transposition. Although the L and R ends feature repetitive TnsB binding sites and are reminiscent of the terminal inverted repeats characteristic of other transposon families, the positioning of these binding sites is distinct on both ends, leading to a striking asymmetry that promotes polarized insertions. Type I-F CASTs generate integration products in both of two possible orientations, referred to as T-LR and T-RL (**Fig. 1D**), but the T-RL products are highly preferred at ratios typically over 90%, depending on the crRNA and CAST system^32,33,41^.

In our early work, we discovered and characterized RNA-guided DNA integration using a representative Type I-F CRISPR-associated transposon from *Vibrio cholerae*, which was assigned the transposon identifier, Tn6677^33,57^. (Although we previously referred to the system as VchINTEGRATE, for insertion of transposable elements by guide RNA-assisted targeting, or VchINT for short, hereafter we refer to system as VchCAST to reconcile prior nomenclature choices in the literature.) Genomic insertions were performed in *E. coli* by co-transforming cells with three separate vectors — pDonor encoding the mini-Tn, pQCascade encoding the TniQ-Cascade complex, and pTnsABC encoding the heteromeric TnsABC transposase — and we reported efficiencies of 40–60% using a 980-bp genetic payload. We also readily achieved integration of DNA payloads up to 10 kb in size and detected integration at 24 genomic target sites tiled across the *E. coli* genome, highlighting the system’s robust programmability. We adopted a high-throughput sequencing approach to unbiasedly query the genome-wide specificity of integration products, termed transposon-insertion sequencing (Tn-seq)^31,33^, and found that VchCAST exhibited remarkable fidelity, with the majority of crRNAs directing >95% on-target accuracy and many exceeding 99%. Notably, similar experiments with Type V-K CAST systems, including ShCAST and ShoCAST (formerly ShoINT), have revealed considerably lower fidelities and Cas12k-independent integration activity^32,42,48^, indicating that some CAST systems retain the ability to undergo random, untargeted transposition.

While our initial study introduced VchCAST as a promising genome engineering platform, the requirement for three separate plasmids encoding multiple expression cassettes complicated construct customization and delivery, limiting broader applications of the system. To address this, we designed a streamlined expression cassette termed pEffector, which expresses all necessary protein and RNA components from a single polycistronic promoter, and then combined this with the mini-Tn genetic payload to generate single-plasmid integration vectors, termed pSPIN^32^ (**Fig. 2**). These vectors considerably simplified CAST delivery to diverse bacteria, led to integration efficiencies approaching 100% after promoter and backbone optimization, and enabled straightforward plasmid curing to remove the CAST system from cells after the desired genomic insertion was introduced^32^. Parallel efforts from Doudna and colleagues similarly highlighted the versatility, efficiency, and specificity of single-plasmid Type I-F CAST systems for bacterial genome engineering, termed VcDART^37^. Interestingly, we found that VchCAST exhibited higher integration efficiencies for large genetic payloads when cells were incubated at temperatures below 37 °C, without a detectable change in genome-wide specificity^32^. Although the molecular basis for this temperature effect is not yet fully understood, it provided an accessible strategy to increase the yield of edited cells with large, ∼10-kb genetic payloads, and may be worth exploring further for certain downstream applications.

**Fig 2.**
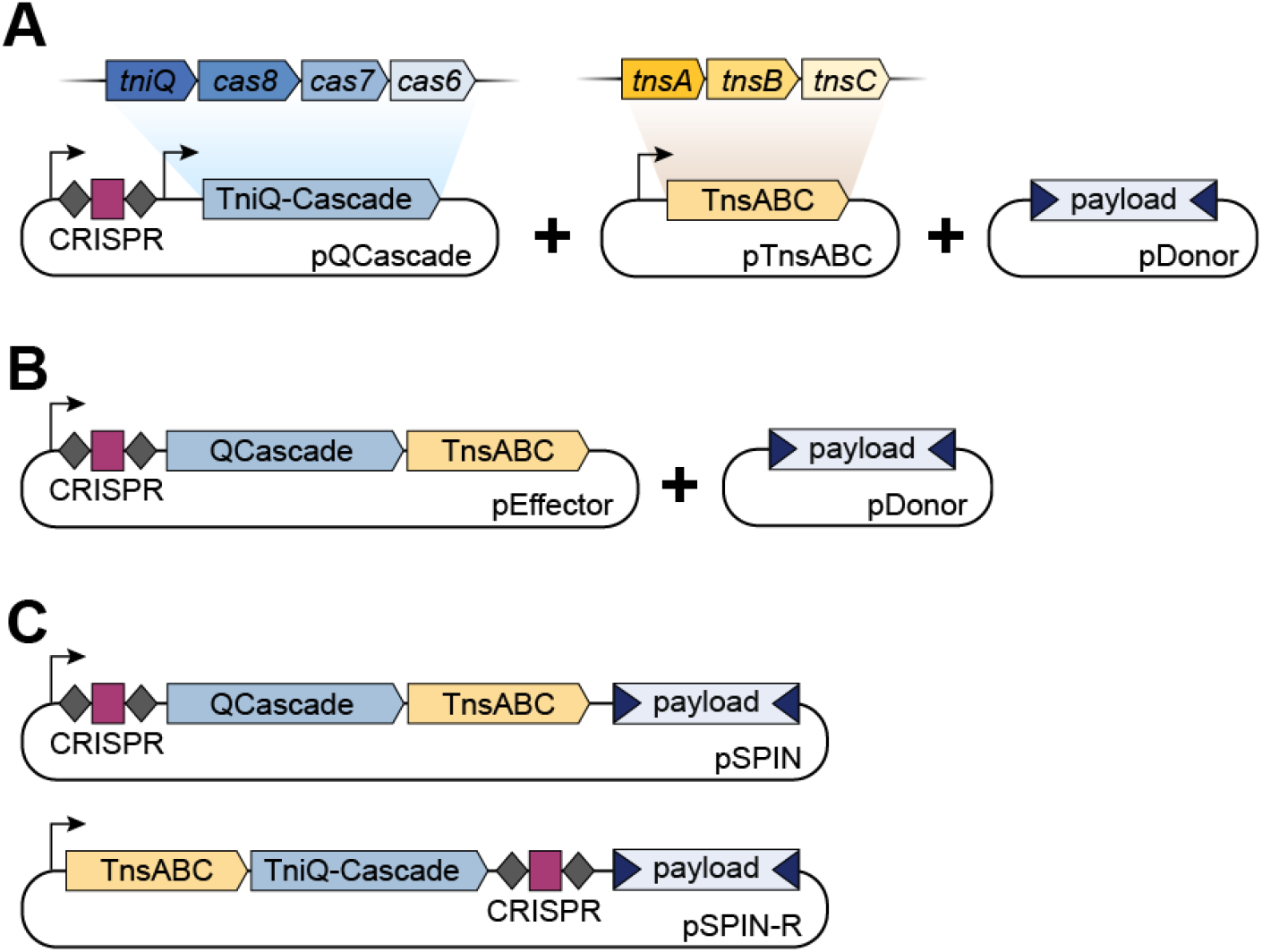
Architecture of VchCAST vector constructs. (A) A 3-plasmid system for CAST expression includes a vector that encodes the crRNA and QCascade components (pQCascade), a vector that encodes TnsABC (pTnsABC), and a final vector that harbors the payload flanked by transposon ends (pDonor). (B) A 2-plasmid system consists of an expression vector (pEffector) vector with single promoter driving expression of crRNA, QCascade, and TnsABC, and separate vector harboring the mini-Tn (pDonor). (C) Highly efficient single-plasmid integration vectors (pSPIN) exist in two variants: (top) pSPIN with a single promoter driving the expression of the crRNA, QCascade, and TnsABC followed by the mini-Tn, and (bottom) pSPIN-R that reduces self-targeting insertions, in which the CRISPR array is relocated just upstream of the mini-Tn.

### Multiplexed insertions

Unlike site-specific transposases and integrases such as Tn7 and Bxb1, which perform efficient genomic integration but cannot be reprogrammed, CAST systems can be easily programmed to insert at new user-defined target sites, and crucially, can be programmed with multiple guide RNAs to insert at multiple user-defined target sites^32,35^. For example, we have shown that multi-spacer CRISPR arrays are efficiently processed into multiple crRNAs in bacteria (**Fig. 3A-B**), leading to multiplexed and simultaneous insertion of the same DNA payload at up to 3 target sites, enabling rapid generation of insertional knockouts^32,35^. Additionally, by encoding a loxP sequence within the mini-Tn payload, we combined VchCAST with Cre recombinase to mediate seamless, programmed deletions of large genomic sequences^32^ (**Fig. 3A**). Separately, Yang and colleagues employed multi-spacer CRISPR arrays alongside single-spacer arrays targeting multi-copy genomic loci, in a strategy that produced *E. coli* strains containing up to 10 genomic insertions of a glucose dehydrogenase expression cassette^35^. Finally, the use of pooled guide RNA libraries with both Type I-F and Type V systems across a population of cells enables efficient disruption of a subset of genes of interest^36,50^, analogous to the use of Cas9 and guide RNA libraries for genetic screening experiments in eukaryotic cells.

**Fig 3.**
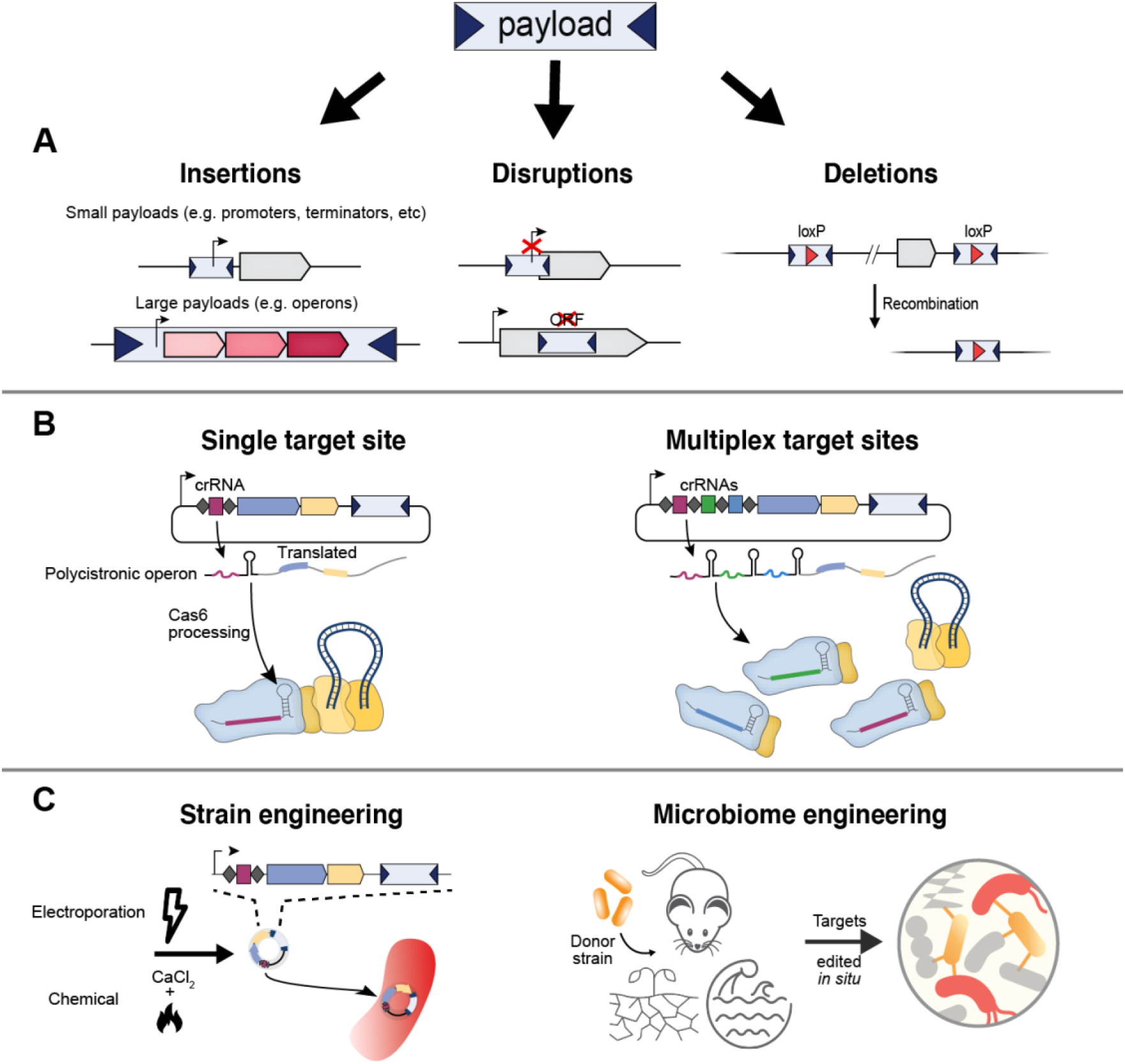
Possible applications of CAST systems for microbial engineering. (A) Schematic depicting the range of genetic modifications that can be generated with CASTs, which include insertions of either small payloads (e.g. promoters, terminators, repressors, etc.) or large payloads (e.g. genes or metabolic operons); gene disruptions via insertion into promoters or gene bodies that disrupt open reading frames (ORF); or programmed deletions via insertion of loxP sites and subsequent Cre-based recombination. Functional insertions can also be generated that simultaneously disrupt endogenous genes. (B) The modifications in (A) can be made at single target sites (left) or multiple target sites using multi-spacer CRISPR arrays (right). crRNAs are first transcribed as a precursor transcript that is processed by Cas6 to form mature crRNAs. (C) RNA-guided insertions can be used to engineer single strain isolates using typical transformation methods, or to precisely modify complex microbial communities in diverse environments (e.g. gut, soil, and/or aquatic microbiomes) via conjugative delivery.

### Homologous and orthogonal Type I-F3 CAST systems

CRISPR-Cas systems are highly diverse, and bioinformatic and experimental mining efforts over the years have repeatedly uncovered new variants that offer advantages for technology development. Thus, we and others have been similarly motivated to explore and develop additional CAST elements for programmable RNA-guided DNA integration^41,43,58^. We recently employed a bioinformatics pipeline to identify hundreds of Type I-F CAST systems in sequenced bacterial genomes, and then experimentally characterized 18 new systems that exhibited integration activity in *E. coli*^41^. These homologs display interesting behaviors when compared to VchCAST, including intriguing modularity between the CRISPR and transposase components, the presence of additional targeting proteins or protein-protein fusions, and unique recognition of transposon end sequences. We identified four additional CAST systems that were capable of highly efficient and accurate DNA insertions, forming a suite of high-value CAST systems for bacterial genome engineering alongside VchCAST. Importantly, these five systems are completely orthogonal: the transposon end sequences from each system are effectively invisible to the transposase machineries from the others, such that genetic payloads encoded within the mini-Tn can be selectively acted upon by only the cognate CAST machinery. This orthogonality could allow consecutive insertions to be generated within a focused genomic region of interest, without the inhibitory consequences of the target immunity pathway that normally precludes tandem insertions at the same site^54^. Collectively, these diverse CAST systems open up the possibility of rapid and iterative engineering of target strains, as well as tailoring system choice for applications within new bacterial species of interest.

### Applications of CAST systems

CAST systems provide a simple yet powerful platform for inserting DNA payloads ranging from hundreds to thousands of base pairs in size (**Fig. 3A**). Smaller payloads can include regulatory elements such as promoters to drive inactive genes, terminators to decouple transcription and translation, and other types of regulatory elements (**Fig. 3A**). Larger payloads, in contrast, can harbor entire operons that enable metabolic rewiring within a target bacterial strain (**Fig. 3A**). As transposons are ubiquitous selfish genetic elements that often mobilize across various hosts, they are particularly promising for applications across diverse bacterial species and strains that have never been genetically modified before (**Fig. 3C**). Although VchCAST was initially characterized in standard *E. coli* strains, we and others have demonstrated its ability to mediate robust genomic integration in multiple strains including recombination-deficient *E, coli* strains^32,33,35,42^, multiple species of *Klebsiella* and *Pseudomonas*^32,37^, as well as in *Tatumella citrea* (**Fig. 3C**) ^35^. In addition, the ability to encode all necessary molecular components on single pSPIN vectors has opened the door to seamless delivery of CAST systems between bacteria via conjugation, the exchange of genetic material through direct cell contact. An exciting application of CASTs in various non-model bacteria is multiplexed targeted gene knock outs, all at one time, using a multi-spacer CRISPR array (**Fig. 3A-B**). This approach can uncover the functions of unknown genes essential to environmental conditions and perturbations, akin to genome-wide transposon mutagenesis libraries but with a more targeted resolution^31^. CASTs thus hold the promise to uncover the dearth of known biological functions that exist in the genomes of bacteria.

In addition to rapidly engineering bacterial species through loss-of-function insertional mutagenesis, the integrative capacity of CAST systems also allows for stable genomic integration of transgenes and potentially entire operonic pathways, with high efficiency and specificity^32^ (**Fig. 3A**). In a demonstration of rapid strain optimization for the biosynthesis of key industrial compounds, Yang and colleagues exploited VchCAST to perform mutliplexed, multi-copy genomic integration of synthesis pathways while also disrupting undesired host degradation pathways^35^ (**Fig. 3B**). This strategy produced modified strains displaying more robust and stable generation of compounds of interest compared to plasmid-based expression.

While engineering individual strains is often desirable (**Fig. 3C**), microorganisms naturally exist within consortia in ecosystems, as members of complex communities with other bacteria, archaea, and eukaryotes. Thus, studying bacteria in isolation limits our understanding of their natural physiology. We and others have developed a foundation to use CAST systems for *in situ* engineering of target species within complex bacterial communities^32,37^, including species that have so far been challenging to edit using existing technologies (**Fig. 3C**). The previously described CAST-based genetic engineering approaches applied to microbial communities can pave a path that bridges the gap between our functional understanding of few cultivable microbes, versus the overwhelming diversity of uncultivated microbes found in all ecosystems. In addition, integrating desired genetic payloads into microorganisms will increase the range of possibilities for probiotic-based therapies. Future applications focusing on broad-host range CAST expression vectors and transient delivery, obviating the need for any stable plasmid maintenance, will vastly expand the toolkit for microbial engineering.

### Comparisons with existing methods

Transposases and integrases are versatile and pervasive genes across all domains of life^59^ and form the foundation for many existing technologies that mediate large DNA insertions into bacterial genomes. For example, systems such as the ICEBs1 integrative element, Cre-loxP recombinase system, or the Tn*7* transposon, have been used for chromosomal integration of exogenous genes and pathways into diverse bacteria^28,60,61^. However, these systems are only able to recognize fixed, system-specific target sequences, and thus, such sequences must either exist in the target strain prior to editing or be installed separately using an orthogonal method^28,60,61^. Other transposon systems such as *mariner* or Tn*5* facilitate genomic insertions with minimal sequence specificity^62–65^, and as a result have been useful for genome-wide transposon mutagenesis or high-throughput integration screens^31,65–68^. However, applications requiring targeted insertions at desired loci cannot effectively make use of such non-specific integrative systems, and clones containing inserts at desired locations must first be identified through additional steps, such as whole-genome sequencing. In this case, platforms with native target programmability such as recombination-based strategies and CAST systems would be more suitable to simplify and accelerate engineering workflows.

Compared to recombineering-based technologies, CAST systems offer several key advantages. Recombineering efficiency is generally low (less than 1 in 10^3^–10^4^)^69^ without selection of a co-integrating selectable marker^70^ or CRISPR–Cas-mediated counter-selection of unedited alleles^17^ and, thus, cannot be easily multiplexed to make simultaneous insertions into the same cell. In contrast, CAST systems efficiently mediate insertions of large multi-kb DNA payloads (near 100%) without requiring any selection for integration events^32,37^, which can be beneficial for scenarios where the use of drug markers is undesired. While the requirement for fixed transposon end sequences flanking the insert of interest does prevent scarless insertions, the absence of a recombination-based process for CAST insertions means that homology arms are never necessary. In contrast, donor DNA molecules used in recombineering require homology arms matching sequences flanking the target insertion site, as well as unique selection marker cassettes (albeit significant advancements have facilitated marker removal^71,72^), which quickly becomes time and labor intensive to generate, especially for multiplexed editing experiments^73^. Thus, for applications such as those involving insertional mutagenesis or intergenic integration of multi-kb payloads, we suggest the use of CAST systems.

### Limitations of CAST systems

Mobilization of the mini-transposon by CAST systems requires recognition and binding of conserved transposon end sequences by the transposase machinery. Thus, any DNA payload of interest must be flanked by these end sequences as part of the entire functional mini-Tn (**Fig. 1**). Engineering applications that require scarless insertions should instead utilize other existing methods. In addition, we previously noted some degree of variability in the precise integration site selected by the transposase during transposition: although insertions predominantly occur 49-bp downstream of the 3’-end of the 32-bp target site, the exact distribution of distances sampled across a population of cells depends on local sequence features that have yet to be fully resolved^33,41^. Additionally, Type I-F CAST systems generate insertions in two possible orientations (**Fig. 1**), although T-RL products are preferred by ratios typically exceeding 100:1, especially for recently reported homologous CAST systems^38,41^.

CAST systems generate insertions at a fixed distance downstream of the genomic site targeted by the guide RNA. Therefore, the target sequence is not disrupted upon integration, such that persistent expression of the enzymatic machinery could in theory lead to repeated insertions at the same site, resulting in multiple tandem transposon copies. However, the integrated transposon inhibits subsequent transposon insertions nearby through a “target immunity” pathway that depends on molecular interactions between TnsB and TnsC^46,49,74^. Target immunity is strongest for insertions directly adjacent to an existing transposon insertion, and gradually decays at further distances, with around 20% of the expected activity restored at a 5 kb distance^32,46^. This effect strongly inhibits tandem transposon insertions, but nevertheless it has been observed that low levels of these undesired products are generated, particularly in scenarios involving strong constitutive expression of the system over extended periods of time^54^. Moreover, we have previously observed self-targeting integration events (**Box 1**), wherein QCascade targets the spacer sequence within the CRISPR array itself during RNA-guided transposition, leading to insertion events within the expression cassette that can inactivate the CAST system^32^. Another alternative integration product that CAST systems generate are cointegrates, which comprise duplicated transposon copies and genomic insertion of the vector backbone (**Box 1**). While rare in Type I-F CASTs, cointegrates are the predominant product in Type V-K CASTs, which lack TnsA. Using Pacbio SMRT long-read sequencing, we demonstrated the WT VchCAST system predominantly generates simple insertions, whereas a D90A mutation in the TnsA active site instead generated integration products to favor cointegrates (>95%)^46^, underscoring the key role of TnsA in the production of single insertions (**Box 1**).

#### Box 1. Alternative integration byproducts.

Although Type I-F CAST systems overwhelmingly generate single-copy simple insertion products^32,33,46^, users should be aware that alternative and/or undesired integration products are also possible with CAST systems (Type I-F/Type V), including off-target insertions (**Fig. Box 1A**), self-inactivating vector insertions (**Fig. Box 1B**), on-target cointegrate products (**Fig. Box 1C**), and tandem on-target insertions (**Fig. Box 1D**)^45,55,56,81^. Off-target insertions can arise through both RNA-dependent and RNA-independent processes, and appear to be much more prevalent (**Fig. Box 1A**) for Type V-K CAST systems (e.g. ShCAST) than for Type I-F CAST systems (e.g. VchCAST), though the underlying molecular basis for this difference is unknown.

**Fig. Box 1.**
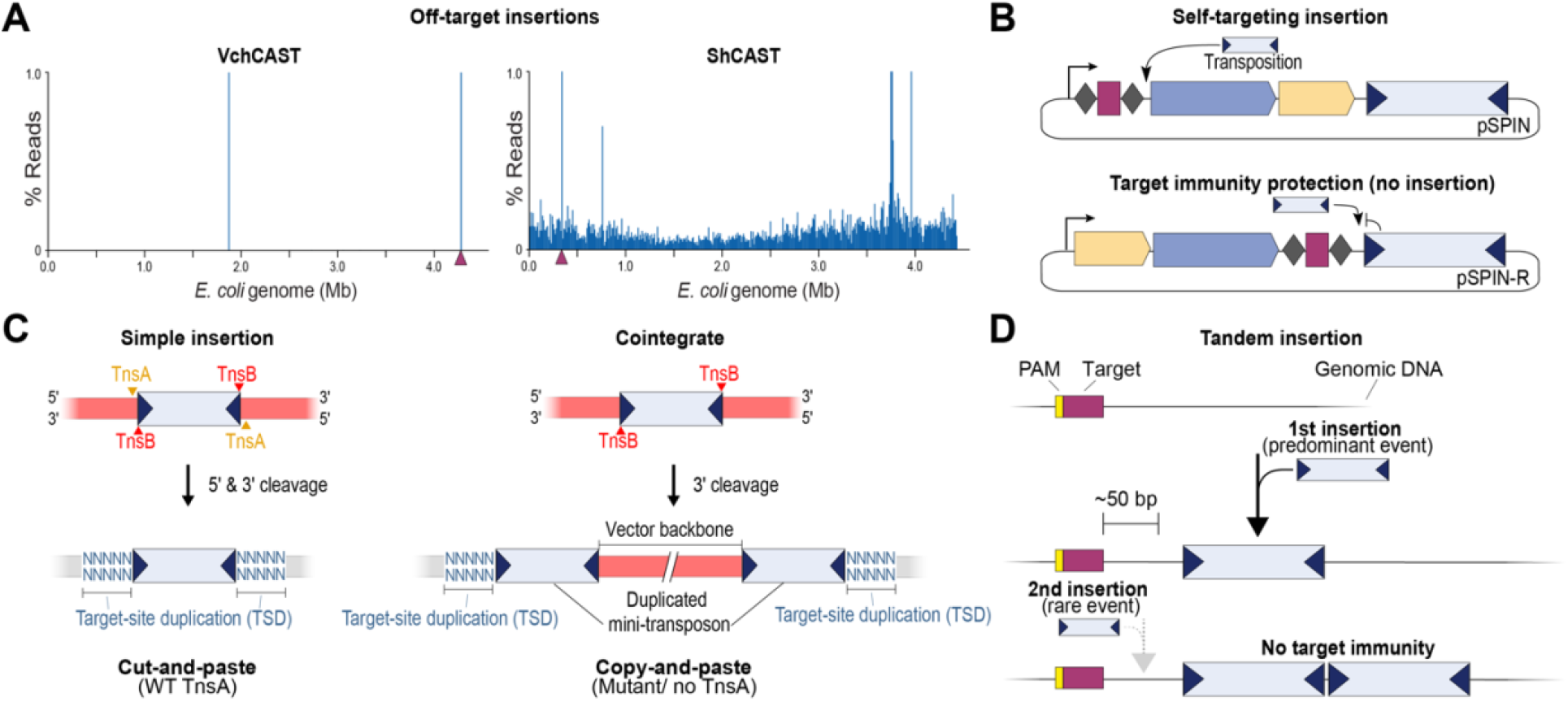
Alternative integration byproducts. (A) Comparison of genome-wide specificity between VchINT (Type I-F) and ShCAST (Type V-K), as assessed via random fragmentation-based NGS library preparation, focused on reads comprising 1% or less of genome-mapping reads. The Type I-F system exhibits exquisite accuracy with low off-target insertions, whereas both Type V-K systems exhibit rampant, off-target integration across the *E. coli* genome. (B) A depiction of undesirable self-targeting insertion events that can occur downstream of the CRISPR spacer due to flexible PAM recognition, and a redesigned vector (pSPIN-R) that ablates self-targeting integration products by harnessing the mechanism of target immunity. (C) Type I-F CASTs can generate simple insertion products via non-replicative, cut-and-paste transposition (left), whereas Type V-K CASTs lacking TnsA (or Type I-F CASTs with inactivated TnsA) generate cointegrate products via replicative, copy-and-paste transposition (right). Cointegrates contain two copies of the inserted mini-Tn that flank the vector backbone. (D) A depiction of low-frequency tandem payload insertion events in Type I-F CASTs.

We previously reported that QCascade-directed DNA integration exhibits a high degree of PAM promiscuity, including low activity with the mutant 5′-AC-3′ self-PAM present within CRISPR repeats flanking the spacer. Thus, low-level targeting of the spacer found within CRISPR arrays can lead to self-targeting insertions that inactivate pSPIN vector, which we detected in the majority of our *E. coli* Tn-seq datasets^32^ (**Fig. Box 1B**). To circumvent this undesired self-targeting product, we redesigned the expression vector such that the CRISPR array is positioned in close proximity to the mini-Tn itself, thereby become largely protected from self-targeting due to the mechanism of transposon target immunity. These modified pSPIN-R vector designs completely abrogated self-targeting, at least for the tested conditions (**Fig. Box 1B**). We therefore encourage users to carefully monitor background levels of vector inactivation due to self-targeting, and to consider use of pSPIN-R vectors instead.

Simple insertion products result from non-replicative, cut-and-paste transposition that relies on the TnsA endonuclease and TnsB transposase for 5’ and 3’-end cleavage at the donor site, respectively, leading to excision of the transposon DNA as a molecule of double-stranded DNA. TnsB subsequently catalyzes strand-transfer reactions at the target site using both 3’ ends of the transposon, followed by gap repair to produce the final integration products flanked by a 5-bp target-site duplication (TSD) (**Fig. Box 1C**). However, inactivating mutations in the TnsA active site result in the formation of so-called cointegrate products, which arise from replication-dependent copy-and-paste transposition and are characterized by the presence of two transposon copies flanking the entire donor vector backbone (**Fig. Box 1C**). Most notably, Type V-K CAST systems lack TnsA altogether and exclusively generate cointegrate products, though these products can resolve to simple insertions through homologous recombination.^45,46,55,56^ We and others have applied PCR-free long-read sequencing to unbiasedly characterize and quantify both types of integration products, demonstrating that Type I-F CAST systems show optimal product purity, but we also encourage interested readers to consider engineered Type V-K CAST systems (HELIX) that show improved properties for both transposition pathway and specificity. Lastly, both types of CAST systems can also generate tandem insertions (**Fig. Box 1D**), since the target site complementary to guide RNA is not destroyed during the integration reaction itself. The frequency of tandem insertions^54^ is naturally limited through the mechanism of target immunity^32,45,46,52,74^, and it can be further reduced by restricting the duration of CAST expression.

### Experimental Design

The generation of programmable genomic insertions using CAST systems involves five main steps (**Fig. 4**): (I) Designing the crRNA and target DNA sequence; (II) cloning the crRNA guide sequence and custom genetic payload into appropriate vectors; (III) delivering one or more vector construct(s) into target cells; (IV) culturing and selection, and (V) analyzing integration events, with optional isolation of desired clones. In the sections below, we provide detailed guidelines for each of these steps to enable use of CAST systems for different engineering scenarios and applications.

**Fig 4.**
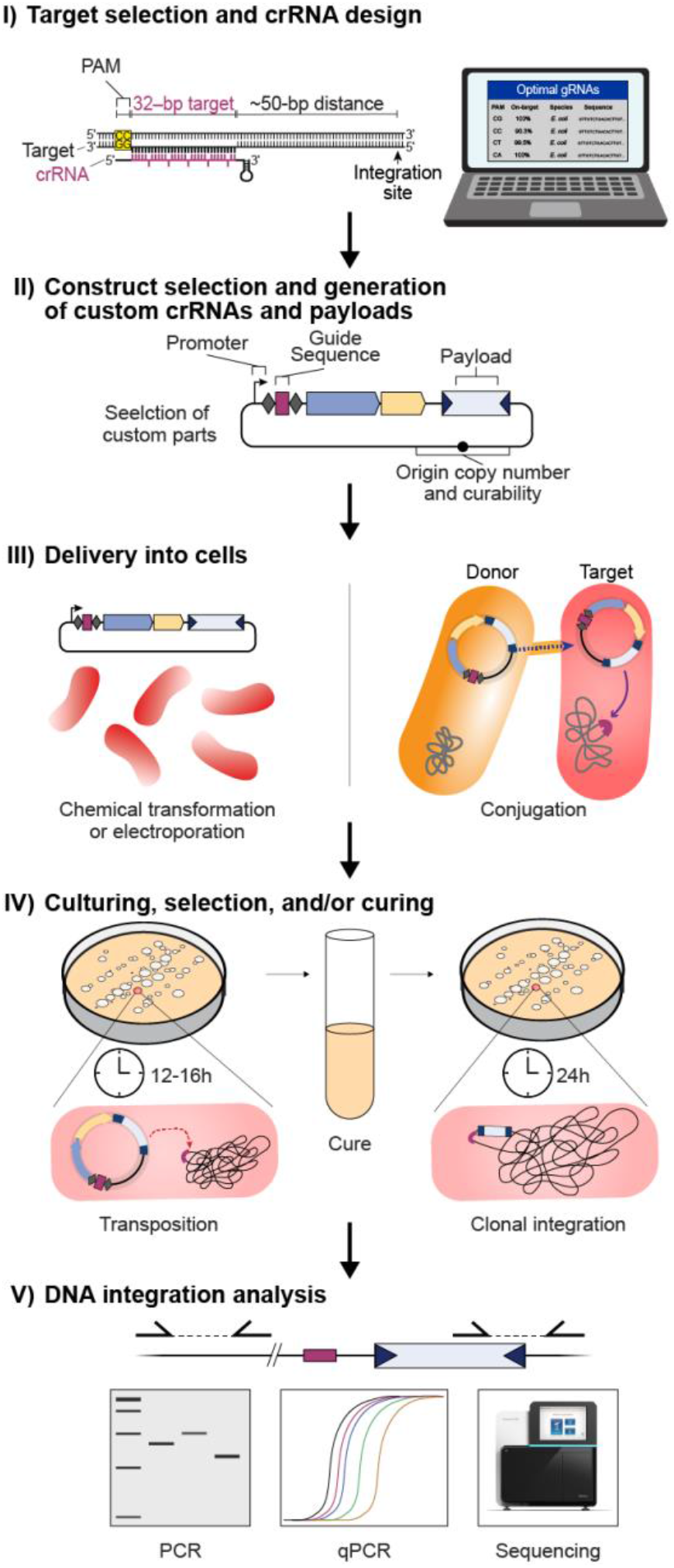
General CAST engineering workflow. (I) The protocol workflow begins with computational analysis of the bacterial genome(s) of interest to identify a 32-bp target site flanked by an appropriate 5’-CN-3’ PAM, which minimizes the likelihood of off-target insertions. (II) One or more appropriate vectors are selected for the desired experiments, and custom crRNAs and DNA payloads (and optional promoters) are cloned. (III) The finalized vector is delivered into target bacteria using either chemical transformation, electroporation, or conjugation. (IV) Target bacteria are incubated for 12–18 h on the appropriate selectable marker to allow efficient transposition to occur, followed by optional curing of the expression plasmid. (V) RNA-guided DNA integration can be assessed using PCR and/or qPCR approaches, and high-throughput sequencing may be used to more systematically evaluate genome-wide specificity during the editing experiment.

### I) Target selection and crRNA design

The general workflow for target selection and crRNA design begins with selecting the desired genomic site for integration of the genetic payload, followed by identifying a 32-bp target sequence located ∼50 bp away. Most insertions occur in a T-RL orientation, with the transposon right end integrated proximal to the target site (**Fig. 1D**), such that insertions in a preferred orientation can be generated by selecting a candidate target site either downstream or upstream of the integration site. The target sequence must be directly flanked by a compatible protospacer adjacent motif (PAM) recognized by QCascade, which is 5’-CN-3’ for most Type I-F CAST systems^33,41,75^. Off-target transposon insertions can occur when genomic sites are highly similar to the intended target site^32^, and the target sequence should therefore be carefully selected such that guide RNAs are avoided with highly similar, partially matching target sites elsewhere in the genome of interest. Mismatches within the seed region (positions 1-8) and the PAM distal region (positions 25-32) are discriminated particularly well by QCascade, whereas mismatches within positions 9-24 are less well discriminated^33,50^, and these factors should be taken into account when evaluating the risk of off-target insertions. However, users may want to perform analyses at candidate off-target sites to confirm the lack of undesired insertions for an isolated clone, and/or perform unbiased off-target insertion profiling.

### II) Construct selection and generation of custom crRNAs and payloads

A suggested set of CAST constructs is provided through Addgene (**Supplementary Table 1**). Single-plasmid pSPIN constructs for the VchCAST type I-F system are encoded on either a pCDF backbone for use in *E. coli*, on a pBBR1 broad-host vector^76^, suitable for use in *E. coli* and related Gram-negative bacteria, on a temperature-sensitive pSC101* backbone that enables simple plasmid curing in *E. coli* cells^32,77^. These constructs should be suitable for many engineering applications. VchCAST may also be delivered to cells as two separate plasmids, in which pEffector encodes the guide RNA and all protein components, and pDonor encodes the mini-transposon payload. These compatible plasmids are provided on *E. coli*-specific vector backbones and can be used in lieu of pSPIN for applications involving large genetic payloads that may be difficult to clone and deliver on a single pSPIN vector (**Fig. 3**). The RNA and protein components are expressed from a strong constitutive promoter (J23119) on most of the vectors we recommend, though this promoter can be easily replaced with alternative promoters that are weaker, inducible, and/or specific to other desired bacterial species. While the overall rates of self-targeting for CASTs are generally low, we also designed a re-engineered variant of pSPIN, termed pSPIN-R, which encodes the CRISPR array proximal to the mini-Tn in order to repress self-targeting through target immunity^32,52^. This construct effectively restricts self-targeting-based vector inactivation but does exhibit slightly lower integration efficiencies compared to pSPIN.

Beginning with pEffector or pSPIN entry plasmids, new crRNA spacer sequences are cloned by ligating hybridized oligonucleotide pairs (outlined in Steps 1-12, **Box 2**). For multiplexed applications, multiple spacers along with intervening repeat sequences can similarly be cloned using several overlapping oligo pairs (**Box 2**). Custom mini-transposon payload sequences can be cloned into pDonor or pSPIN vectors through various simple cloning strategies, thought we outline steps for Gibson assembly within this protocol (Step 13). We have shown robust integration activity with transposons ranging from ∼300 bp to ∼10 kb in size^32^. As natural CRISPR-associated transposons can be as large as 100 kb in length^34,43^, payloads much larger than 10 kb could in theory be mobilized, however, the full size range has not yet been systematically investigated. In general, peak integration efficiencies in *E. coli* under certain experimental conditions were observed with mini-Tn constructs spanning 500-1000 bp in size, and larger and smaller transposons may result in decreased efficiencies.

#### Box 2. Cloning CRISPR spacers for CAST systems.

The first step in CAST guide RNA design is to identify available PAMs that would place the desired transposon integration site 48-50 bp downstream of the adjacent 32-bp target (or 70– 72 bp downstream of the PAM). The VchCAST systems (Tn6677) recognizes 5’-CN-3’ PAMs, whereas other homologous Type I-F CAST systems are even more permissive^41^, thus offering a flexible targeting window. CRISPR arrays encoding the candidate crRNA guides should be designed so that the 32-bp spacer sequence exactly matches the 32-bp immediately 3′ of the PAM on the target genome (**Fig. 1D**). crRNA spacers with a high degree of sequence identity to other regions of the genome should be filtered out, in order to minimize the risk of off-target insertions. We encourage users to use an in-house pipeline we developed to screen candidate crRNA using a BLAST-based approach, which is available on Github at https://github.com/sternberglab/CAST-guide-RNA-tool.

Once final spacer sequences have been chosen, they are cloned via ligation into pEffector or pSPIN entry vectors, in which Type II-S (BsaI or BbsI) restriction sites are flanked by two CRISPR repeats (**Fig. Box 2B**). A pair of oligonucleotides should be designed, in which oligo-1 contains 5-nt of overlap with the plasmid digestion site at the 5’ end, followed by the 32-nt spacer sequence, and ending in 1-nt overlap with the plasmid digestion site at the 3’ end. Oligo-2 contains the reverse 32-nt spacer, again with a 5-nt overlap at the 5’ end and a 1-nt overlap at the 3’ end, as shown in **Fig. Box 2B**. Both oligos are hybridized together to form a sticky-end DNA product containing the crRNA spacer of interest, for ligation into pEffector or pSPIN that was digested with BsaI or BbsI (depending on the vector).

**Fig. Box 2.**
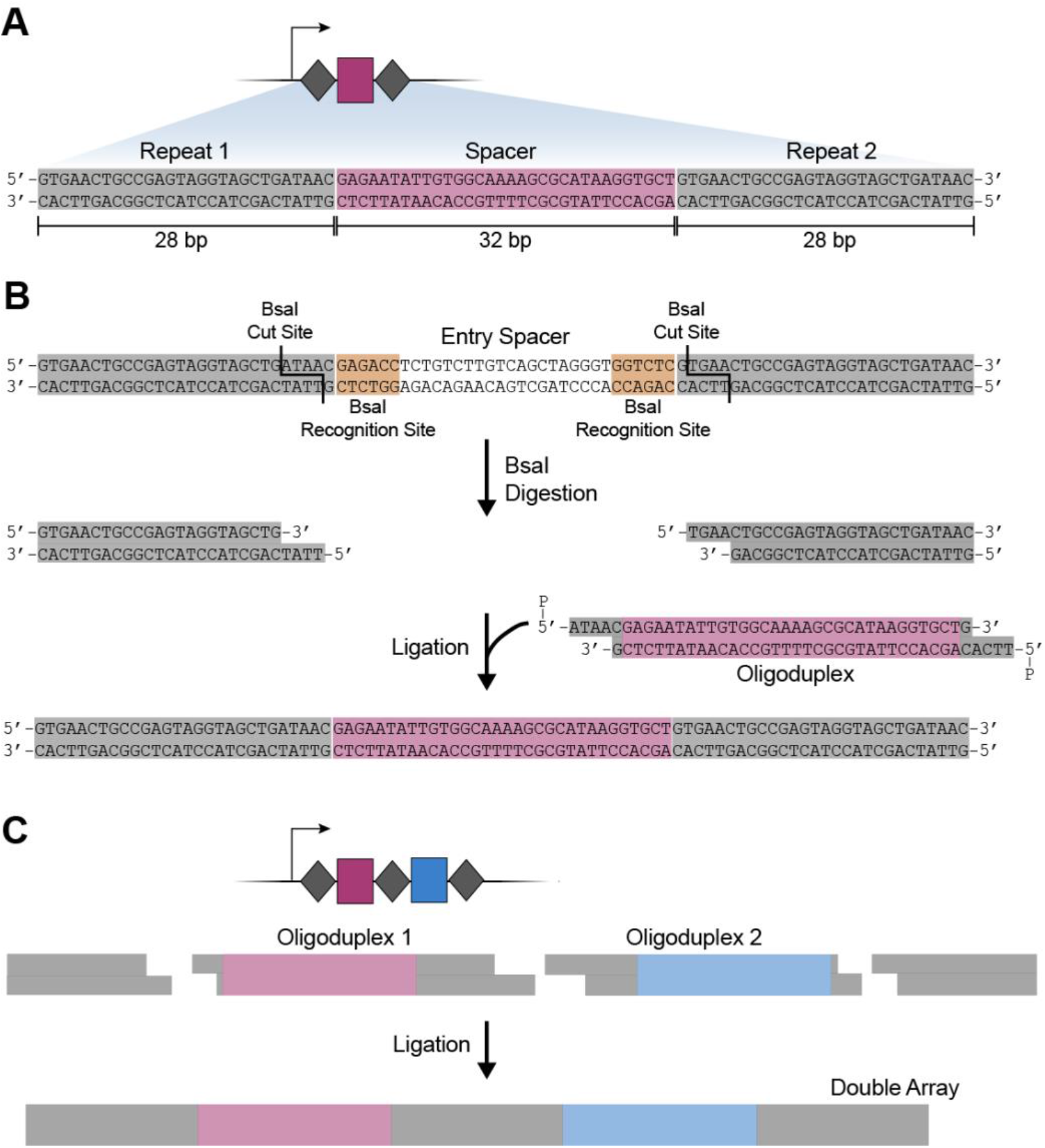
Cloning CRISPR spacers for CAST systems. (A) Diagram of a single-spacer CRISPR array for VchCAST, characterized by two 28-bp repeats that flank a 32-bp spacer complementary to the target site. (B) Schematic outlining the strategy to clone spacers into expression vectors using BsaI restriction digestion and oligoduplex ligation. (C) Schematic for a cloning strategy to generate multi-spacer CRISPR arrays, for multiplexed RNA-guided DNA insertions.

Multiplexed CRISPR arrays, which encode distinct crRNAs to enable targeted insertion at multiple genomic sites, can also be constructed following a similar design and cloning strategy. The main difference is that rather than using a single pair of hybridized oligos, multiple oligoduplexes are combined into a single ligation reaction that yields the desired CRISPR array (**Fig. Box 2C**).

A recent report described the presence of a promoter within non-essential regions of the VchCAST transposon right end^33,35^, which can lead to leaky expression of encoded payloads genes. In addition, we have observed improvements in the product purity and efficiency using transposon variants containing truncated right ends missing this region^33^. Thus, for all constructs in this protocol, we encourage use of modified VchCAST mini-Tn designs in which the right end is truncated to a final length of 57-bp relative to our initial reports^32,33^. Similar leaky expression may also occur with other homologous Type I-F CAST systems, and we point interested readers to recent work that pursued a more systematic investigation of the minimal left and right end sequence requirements during VchCAST transposition^78^.

### III) Delivery into cells

Within the context of common *E. coli* lab strains, simple heat-shock transformation of chemically competent cell preparations is sufficient for vector delivery. For other strains and species, we suggest high-efficiency electroporation as the default strategy for cellular transformation, particularly for experiments involving large plasmids and/or the combined use of pEffector and pDonor vectors. Of note, even with a low transformation efficiency, the high integration efficiency with CAST systems enables straightforward isolation of clones containing the desired genomic insertion.

For species or strains where electroporation is inefficient or impractical, and especially within complex bacterial communities^32,37,79^, we encourage the use of conjugation as an alternative route for transformation^80^. Bacteria naturally exchange plasmids through conjugation as a highly effective means to share genetic material. We and others have utilized bacterial conjugation to efficiently deliver plasmids into isolates as well as complex bacterial communities, demonstrating its viability for CAST-mediated genome engineering^32,37^. We especially recommend the RP4 transfer system to conjugatively transfer DNA from *E. coli* to several bacterial species due to its broad-range capabilities. In this protocol, we describe steps to generate and transform chemically and electrocompetent *E. coli*, as well as a protocol for transforming *E. coli* through conjugation.

### IV) Culturing, selection, and/or curing

After transformation and recovery, cells are usually plated on solid media with appropriate antibiotic selection. While integration can also be performed within a liquid culture, we have observed that solid plating during transposition reduces potential competitive growth effects within a heterogeneous cell population^32^. As the efficiencies of integration are generally high, we normally select only for antibiotic resistance encoded on the vector backbone(s). (Note that many of the mini-transposon variants in our entry vectors encode a promoter-less chloramphenicol resistance gene that we originally selected as an arbitrary payload construct.) For scenarios where selection for integration events is desired, users should clone a drug marker expression cassette into the transposon. However, as vector-based expression of the transposon marker can also occur under conditions where the vector is stably maintained, users should only perform selection after curing cells of the plasmid. Alternatively, a “promoter capture” approach can be used, whereby a transposon encoding a promoter-less marker gene is inserted downstream of an active genomic promoter. Here, users should take steps to ensure that there is no leaky vector-based expression, such as by truncating the transposon right end (discussed above).

Culturing of *E. coli* cells to allow transposition to occur is typically performed at 37 °C over a period of ∼18-24 h, unless using the temperature-sensitive pSC101 vector. However, we have observed that longer incubation times at either 30 °C or 25 °C can strongly enhance integration efficiencies with VchCAST in *E. coli*, particularly for large DNA payloads^32^. This effect is not universal across other homologous Type I-F CAST systems, nor has it been tested in bacteria other than *E. coli*; however, altered temperature incubations can easily be performed in parallel while optimizing the system for a new target species, particularly for species that do not grow optimally at 37 °C. Multiple cycles of solid-media culturing has also been shown to induce higher efficiencies for multiplexed integrations^32^, and dilution of cultures at early log-phase may also enhance efficiencies when performing integration assays in liquid media^32^.

If using pSPIN constructs on the temperature-sensitive pSC101 vector, cells can be cured of the plasmid after integration via liquid-media growth at 37 °C in the absence of drug selection. Other backbones such as pBBR1 can also be cured by culturing cells over several generations without drug selection, combined with frequent phenotyping on selective media, though with less robustness. While not described in this protocol, users may also explore other published methods for plasmid curing, such as the Cas9-based pCutamp system^81^.

### V. DNA integration analysis

Transposon insertions at the target site can be routinely detected by genotyping using targeted PCR^32,33^. A standard PCR strategy probes for the existence of genome-transposon junctions at the target locus, using a genome-specific primer paird with a transposon-specific primer (**Fig. 5**). Since the VchCAST system produces a low level of T-LR insertions, multiple primers can be designed to probe for both of the two possible orientations, if desired. For experiments where the efficiency of integration is of interest, qPCR can be used to quantify the proportion of genomic DNA molecules containing these junctions^32,33^. We also describe in this protocol a simple PCR strategy involving two genome-specific primers flanking the transposon insertion, which is useful for isolating clonal integrants.

**Fig 5.**
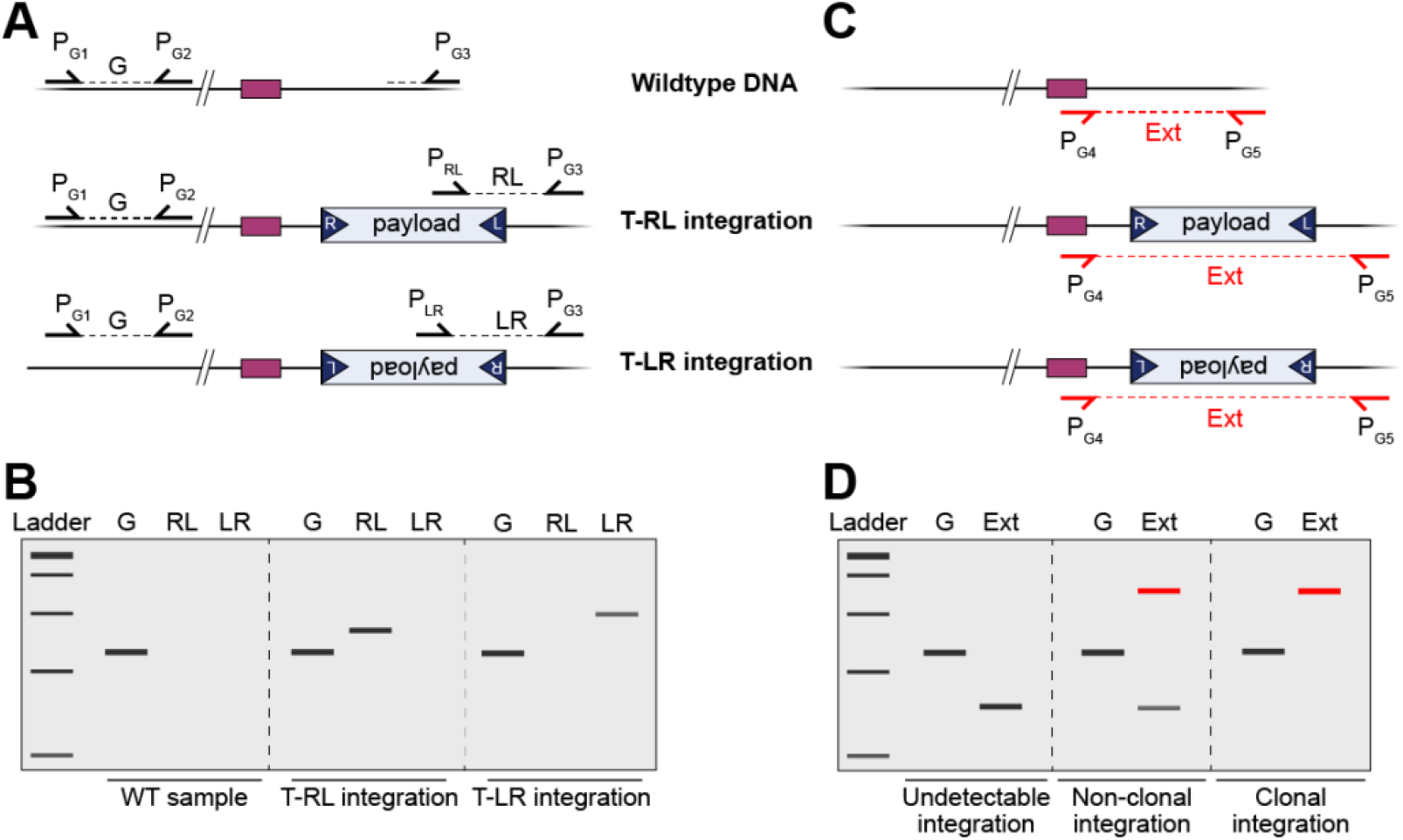
PCR and qPCR analysis of integration products. (A) A schematic representation of junction primer pairs used for the analysis of integration events for unmodified, wild-type DNA (top), T-RL integration products (middle), and T-LR integration products (bottom). PCR product G, amplified by primer pair P_G1_ and P_G2_, refers to a reference gene that is unaffected by CAST targeting and serves as a benchmark for PCR and qPCR measurements. Integration products are detected by priming off the transposon and a genomic site downstream of the integration site, resulting in either junction PCR product RL (primer pair P_RL_ and P_G3_) or junction PCR product LR (P_LR_ and P_G3_). (B) A schematic representation of a gel with the simulated PCR products from (A). Note that individual colonies may also show bands indicative of both the T-RL and T-LR products, due to the presence of heterogeneous alleles present within single (non-clonal) colonies. (C) Integration products can also be detected using primers that bind upstream and downstream of the integration site, rather than the junction strategy shown in (A). The primers (P_G4_ and P_G5_) will generate a PCR product for both the unmodified (WT) DNA and both integration products, with the product size identifying the presence of the desired edit. (D) A schematic representation of a gel with the simulated external PCR products from (C). Note that this method is less sensitive than junction PCR, and that colonies may yield bands for both of the potential products (unmodified and integrated) due to non-clonality.

Importantly, we have repeatedly observed that *E. coli* colonies are often genetically heterogeneous and non-clonal after a single night of culturing and drug selection^33^. Diagnostic PCR analyses demonstrate the concurrent presence of all possible products — no integration, T-RL integration, and T-LR integration — indicating that transposon insertion is slower than the cell doubling time and thus multiple alleles are propagated within the same colony. We therefore encourage users interested in isolating clonal integrants to re-plate cells at least once, in order to allow integration products to be homogeneously fixed within the population of cells in the colony.

## MATERIALS

### Biological Materials

- NEB Turbo (NEB cat. no. C2984), NEB 10-beta (NEB cat. no. C3019), or NEB Stable (NEB cat. no. C3040) *E. coli* cells for cloning (chemically competent or electrocompetent)
- Target bacterial strain-of-interest
  - E.g., *E. coli* BL21(DE3) Chemically Competent Cells (Sigma cat. no. CMC0014)
  - E.g., *E. coli* BL21(DE3) Electrocompetent Cells (Sigma cat. no. CMC0016)
  - E.g., *E. coli* MG1655 (ATCC, cat. no. 700926)
- Conjugative donor strain *EcGT2* (ATCC cat. no. 47055)

### Reagents

#### Plasmids

- Addgene plasmid list (see **Supplementary Table 1**)

#### Common Reagents

- Spectinomycin dihydrochloride pentahydrate (Gold Biotechnology cat. no. S-140-5)
- Kanamycin monosulfate (Gold Biotechnology cat. no. K-120-5)
- Carbenicillin disodium (Gold Biotechnology cat. no. C-103-5)
- LB medium
- LB agar medium
- Agarose, low EEO, molecular biology grade (Fisher, Scientific cat. no. BP160-500)
- TAE buffer, 50X (Bio-Rad, cat. no. 1610773)
- Gel loading dye, 6X (included with enzyme, or purchased separately: NEB cat. no. B7024S)
- SYBR Safe DNA gel stain (Thermo Fisher Scientific cat. no. S33102)
- 1 kb DNA Ladder (Gold Biotechnology cat. no. D010-500)
- 100 bp Plus DNA Ladder (Gold Biotechnology cat. no. D003-500)
- Milli-Q (MQ) water
- Liquid nitrogen
- Absolute ethanol
- Isopropanol

#### Competent cell preparation

- Glycerol
- DMSO
- MgCl_2_ hexahydrate powder
- CaCl_2_ dihydrate powder

#### Conjugation

- Square plates
- 6-well plates
- 1x PBS pH 7.4 (Gibco cat. no. 10010023, or home-made)

#### DNA Extraction Kits

- QIAprep Spin Miniprep Kit (Qiagen cat. no. 27115)
- QIAquick Gel Extraction Kit (Qiagen cat. no. 28706X4)
- MinElute Gel Extraction Kit (Qiagen cat. no. 28604)
- Wizard Genomic DNA Purification Kit (Promega cat. no. A1125)

#### PCR and qPCR

- Q5 Hot Start High-Fidelity DNA Polymerase (NEB, cat. no. M0493S/L)
- dNTP mix, 10 mM (NEB, cat. no. N0447S/L)
- Q5 reaction buffer, 5X (included with enzyme, or purchased separately: NEB cat. no. B9027S)
- (Optional) OneTaq Quick-Load 2X Master Mix (NEB cat. No. M0486S/L), for genotyping only
- Oligonucleotide primers for PCR, from IDT or preferred vendor
- SsoAdvanced Universal SYBR Green Supermix (Bio-Rad cat. no. 1725270–1725275)
- Oligonucleotide primers for qPCR, from IDT or preferred vendor

#### Construct Cloning

- T4 DNA Ligase (NEB cat. no. M0202S/T/L/M)
- T4 Polynucleotide Kinase (NEB cat. no. M0201S/L)
- NEBuilder HiFi DNA Assembly Master Mix (NEB cat. no. E2621S/L/X)
- BsaI-HFv2 restriction enzyme (NEB cat. no. R3733S/L)
- BamHI-HF restriction enzyme (NEB cat. no. R3136S/L/T/M)
- HindIII-HF restriction enzyme (NEB cat. no. R3104T/M)
- KpnI-HF restriction enzyme (NEB cat. no. R3142S/L/M)
- PstI-HF restriction enzyme (NEB cat. no. R3140S/L/T/M)
- XhoI restriction enzyme (NEB cat. no. R0146S/M)
- Bsu36I restriction enzyme (NEB cat. no. R0524S/L)
- SalI restriction enzyme (NEB cat. no. R0138S/T/L/M)
- CutSmart or rCutSmart buffer, 5X (included with enzyme, or purchased separately: NEB cat. no. B7204S/B6004S)
- T4 DNA Ligase reaction buffer, 10X (included with enzyme, or purchased separately: NEB cat. no. B0202S)

#### Equipment

- Glassware
  - Electroporation cuvettes, 1 or 2 mm gap
  - Sterile glass plating beads
  - Assorted glass bottles
- Plasticware
  - Single PCR tubes with attached flat caps, 0.2 mL
  - 8-Strip PCR tubes with attached flat caps, 0.2 mL
  - 96-well PCR plate
  - PCR plate foil seal
  - Conical polypropylene centrifuge tubes, 50 ml
  - Two-sided disposable polystyrene plastic cuvettes, 1.5–2 mL
  - Microcentrifuge tubes, 1.7 mL
  - Sterile filtered pipette tips
  - Sterile pipette tips
  - Sterile 100 x 15 mm petri dishes
  - Sterile plastic inoculation loop
  - Hard-Shell 384-Well qPCR plates, clear shell/white wells
  - Microseal ’B’ qPCR plate sealing film
  - Serological pipettes, 2 mL to 25 mL
  - Sterile baffled plastic Erlenmeyer flask, 250 mL or 2 L
  - Cryotubes
  - Vacuum Filter/storage bottle system, 0.22 µm Pore 33.2 cm^2^ PES Membrane, Sterile (Corning cat. no. 431097)
- Tools and instruments
  - Static incubator
  - Shaking incubator
  - Tabletop centrifuge
  - Benchtop microcentrifuge
  - Benchtop minicentrifuge
  - Microvolume spectrophotometer
  - 96-well thermocycler
  - CFX384 384-well qPCR system
  - Gel electrophoresis power supply
  - Gel electrophoresis tanks
  - Gel casting trays
  - Gel imaging system
  - GenePulser bacterial electroporation system
  - Heat block for microcentrifuge tubes
  - Cell culture spectrophotometer
  - Benchtop vortexer
  - Blue-light gel platform
  - Metal razors
  - Pipets: p1000, p200, p20, p2
  - 8-well multi-channel pipets: p1000, p200, p20, p2
  - Pipet tips: p1000, p200, p20, p2
- Software
  - Benchling, Geneious, or preferred comparable software
  - CAST crRNA design tool: https://github.com/sternberglab/CAST-guide-RNA-tool

#### Reagent Setup

- 50% (v/v) glycerol solution
  - Mix 200 mL of glycerol with 200 mL of MQ water. Sterilize by autoclaving or sterile filtration. Cool down on ice before use; store at 4 °C for < 6 months.
- 10% (v/v) glycerol solution
  - Mix 50 mL of glycerol with 450 mL of MQ water. Sterilize by autoclaving or sterile filtration. Cool down on ice before use; store at 4 °C for < 6 months.
- 1 M MgCl_2_ solution
  - Dissolve 101.65 g of MgCl_2_ hexahydrate powder with MQ water in 250 mL. Once dissolved bring final volume to 500 mL with MQ water. Sterilize by autoclaving or sterile filtration. Store at room temperature.
- 1 M CaCl_2_ solution
  - Dissolve 73.51 g of CaCl_2_ dihydrate powder with MQ water to reach a final volume of 500 mL. Sterilize by autoclaving or sterile filtration. Store at room temperature.
- 100 mg/mL spectinomycin solution (1000X stock)
  - Dissolve 5.0 g of spectinomycin powder with MQ water to reach a final volume of 50 mL, and sterile filter. Aliquot into microcentrifuge tubes and store at -20 °C for < 12 months.
- 50 mg/mL kanamycin solution (1000X stock)
  - Dissolve 2.5 g of kanamycin powder with MQ water to reach a final volume of 50 mL, and sterile filter. Aliquot into microcentrifuge tubes and store at -20 °C for < 12 months.
- 50 mg/mL carbenicillin solution (1000X stock)
  - Dissolve 2.5 g of carbenicillin powder with MQ water to reach a final volume of 50 mL, and sterile filter. Aliquot into microcentrifuge tubes and store at -20 °C for < 12 months. Ampicillin may also be used in lieu of carbenicillin.
- 50 mg/mL diaminopimelic acid (DAP) solution (1000X stock)
  - Dissolve 2.5 g of DAP powder with MQ water to reach a final volume of 50 mL, and sterile filter. Aliquot into microcentrifuge tubes and store at -20 °C for < 12 months.

## PROCEDURE

### Construct selection and crRNA design - [Timing - 3 h]

1) Chose an appropriate CAST vector construct or combinations of constructs (see “Experimental Design - Target selection and crRNA design”). For most use cases, we recommend starting with the VchCAST (Tn*6677*) constructs, specifically the pSC101*-pSPIN vector for *E. coli* applications that require efficient plasmid curing, or the pBBR1-pSPIN vector for integration experiments int *E. coli*, *Klebsiella*, *Pseudomonas*, or related Gram-negative bacteria.
  - Use crRNA tool to design oligos as in **Box 2**

### Cloning custom crRNA spacers - [Timing - 2 d]

2) Digest pEffector or pSPIN in the following reaction mix:

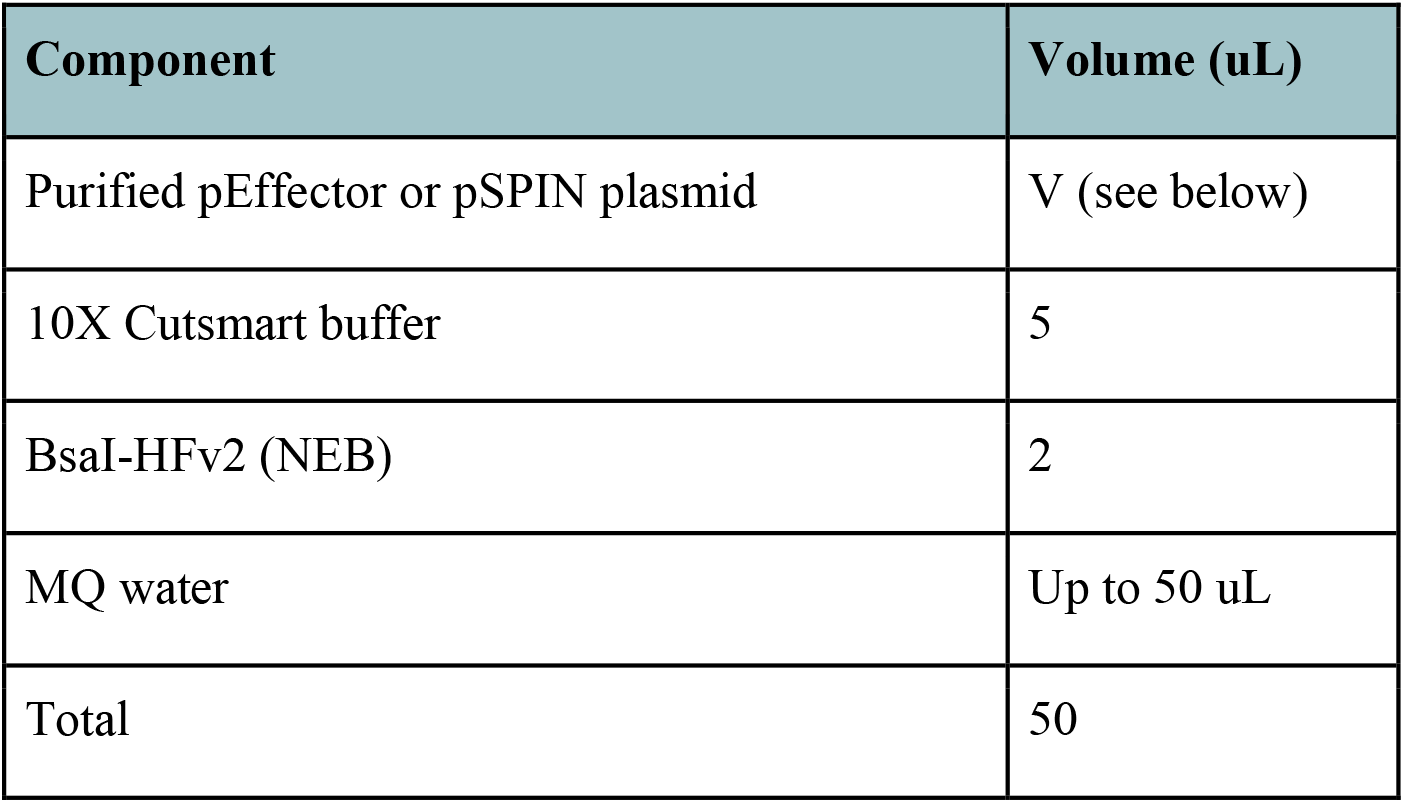 Suggested input DNA amounts:

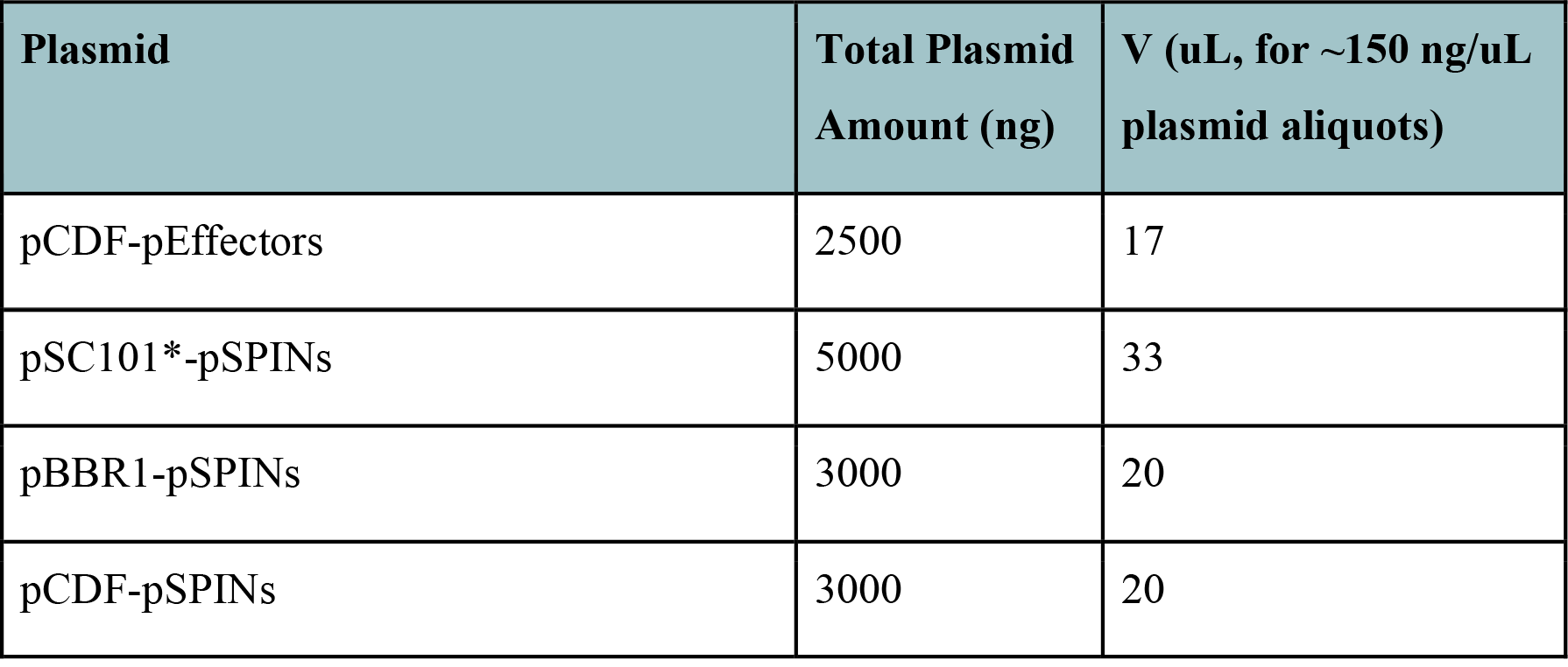 Incubate at 37 °C for 2 h.
3) Add gel loading dye and run the digestion on a 1% agarose gel supplemented with SYBR Safe or another appropriate DNA staining reagent. Typical gel electrophoresis conditions are 140 V for 30 min in 1X TAE buffer.
4) Use a clean blade or razor to cut the band at the expected size (∼12-14 kb, depending on plasmid construct), and extract digested vector using the QIAquick Gel Extraction Kit (Qiagen) according to the manufacturer’s guidelines. If the digested band is faint, the MinElute Gel Extraction Kit (Qiagen) should be used instead to obtain sufficiently concentrated DNA. Elute in 30 uL of EB buffer for QIAquick or 10 uL of EB buffer for MinElute. [PAUSE POINT]
5) During digestion/gel electrophoresis, prepare hybridized oligoduplex for ligation. If primers are purchased from a vendor containing 5’-phosphorylation modifications, mix 2.5 uL of each primer and 20 uL of MQ water and skip to step 5b.
  A. Prepare reactions to 5’-phosphorylate both oliognucleotides, as follows:

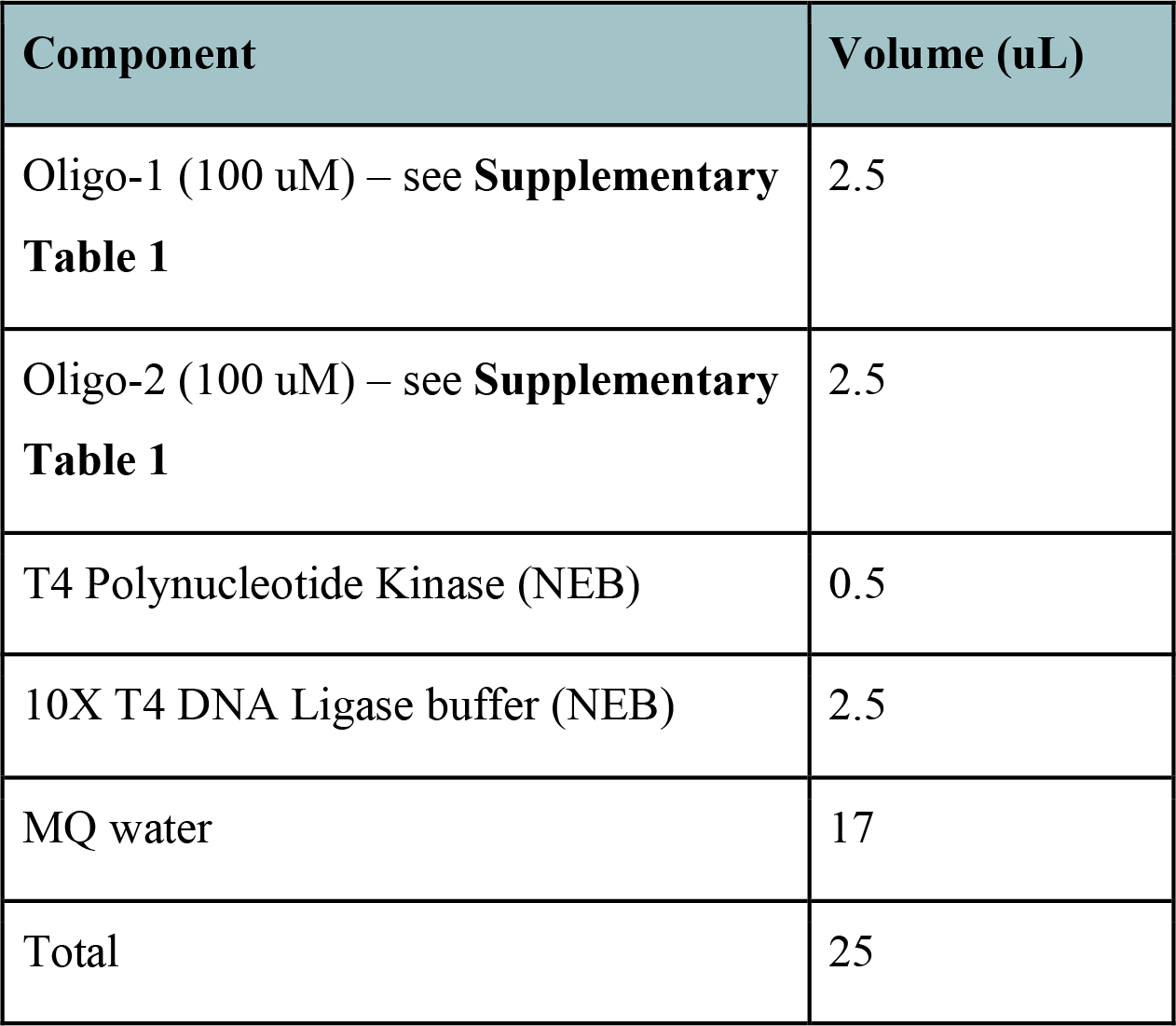 Incubate at 37 °C for 30 min and then heat inactivate at 65 °C for 20 min.
  B. Heat the mixture to 95 °C for 2 min, then allow to gradually cool to room temperature.
6) If ligating a single oligoduplex (i.e. encoding a single spacer), prepare a 1:200 dilution of the oligoduplex mixture into MQ water (50 nM final concentration), then add to a ligation reaction as outlined below. [CRITICAL] Prepare the ligation reaction on ice or add the ligase last, to prevent high levels of spurious intramolecular ligation of the digested vector.

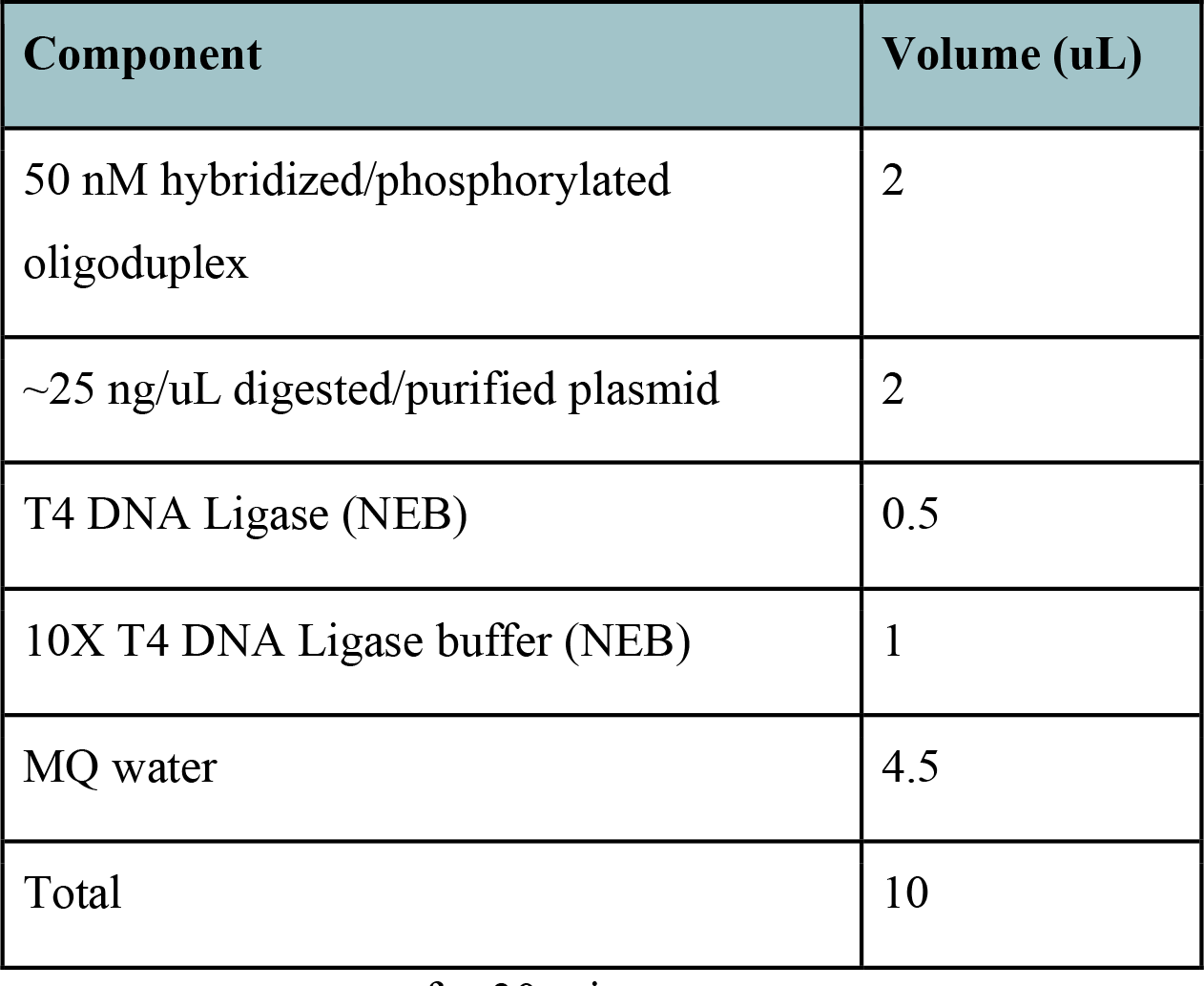 Incubate at room temperature for 30 min. If ligating multiple oligoduplexes (i.e. encoding multiple spacers; **Supplementary Table 2**), prepare a 1:50 dilution of each oligoduplex mixture into MQ water (50 nM final concentration), then add to a ligation reaction as outlined below. [CRITICAL] Prepare the ligation reaction on ice or add the ligase last, to prevent high levels of spurious intramolecular ligation of the digested vector.

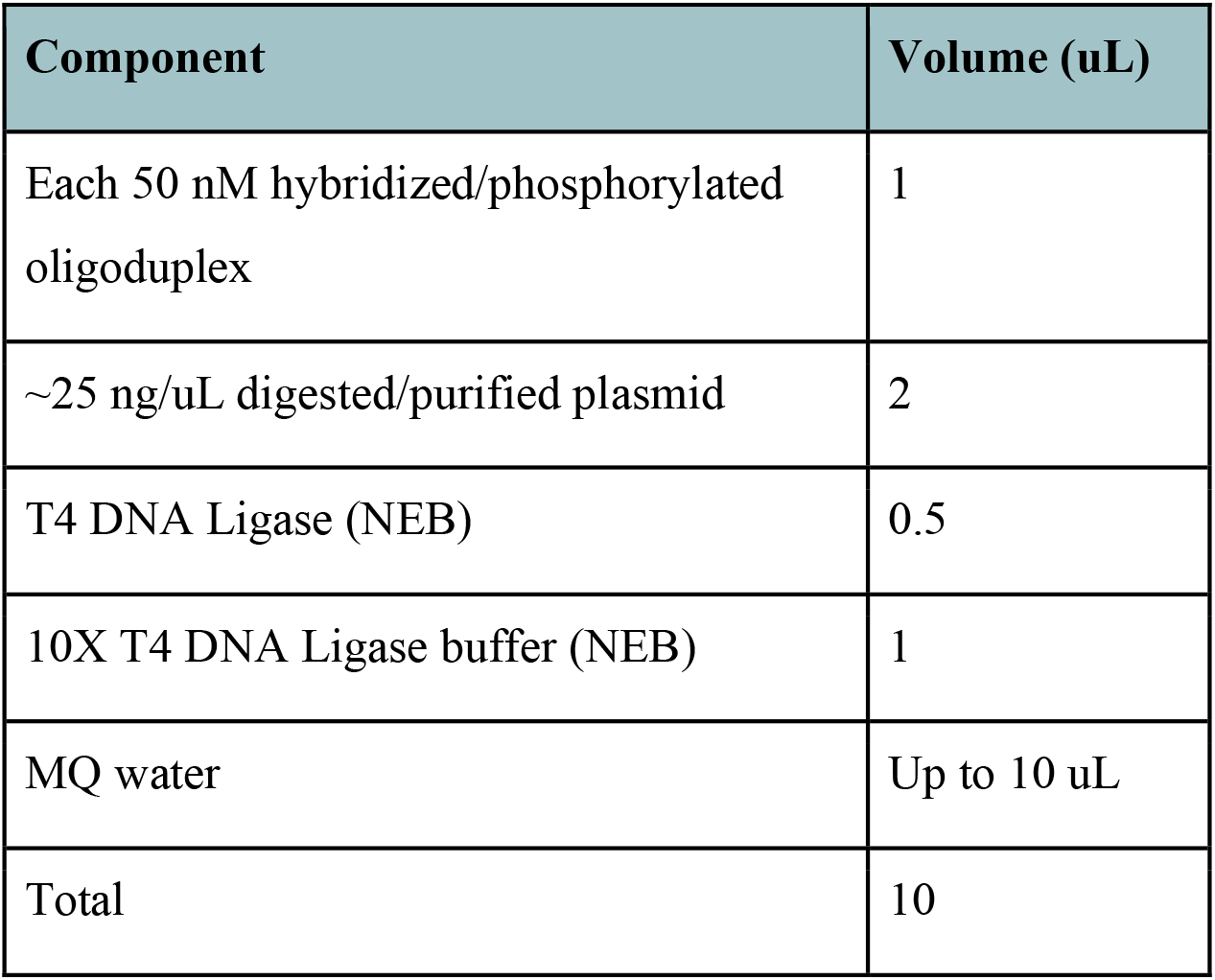 Incubate at room temperature for 30 min.
7) Mix entire ligation reaction with 50 uL of chemically competent *E. coli* cells (cloning strain) on ice. Incubate on ice for 15 min, heat shock at 42 °C for 30 s, then place back on ice for 5 min. [CRITICAL] Commercial, chemically competent *E. coli* (NEB 10-beta, NEB Turbo, NEB Stable, etc.) are recommended.
8) Transfer entire transformation mix to a microcentrifuge tube with 950 uL LB media. Recover with shaking or rotation at 37 °C for 1 h. [CRITICAL] If cloning into pSC101* plasmids, recover at 30 °C for 2 h.
9) Plate 100 uL of recovery on an LB-agar plate containing the appropriate antibiotic (**Supplementary Table 1**). Pellet the remaining 900 uL recovery, discard supernatant, resuspend in 100 uL LB, and plate on a similar LB-agar plate. Ensure that all visible liquid pools have dried off from plates, and incubate plates at 37 °C overnight until colonies are visible. [CRITICAL] If cloning into pSC101* plasmids, incubate cells at 30 °C overnight. Plates may take up to 24 h to produce visible colonies. [PAUSE POINT] [TROUBLESHOOTING] No colonies or high background, see **Supplementary Table 4**.
10) Using 3-6 colonies from the plates in step 9, inoculate a 5 mL culture of LB media containing 1X concentration of the appropriate antibiotic in a 50 mL conical tube. Incubate overnight with shaking at 37 °C. [CRITICAL] If cloning into pSC101* plasmids, incubate cells at 30 °C overnight.
11) (Optional) Cell cultures can be stored at -80 °C for several years as a glycerol stock by mixing 300-500 uL of turbid overnight culture with an equal volume of sterile 50% glycerol. If more plasmid is needed, inoculate LB media as above using a small scraping of this frozen stock; alternatively, transform a cloning *E. coli* strain with 5-10 ng of miniprepped plasmid and inoculate LB media using colonies retrieved from this transformation.
12) Extract plasmid DNA from each culture using QIAprep Miniprep Kit (Qiagen) or similar plasmid extraction kit, and verify the CRISPR array sequence with Sanger sequencing; suggested primers are listed in **Supplementary Table 3**. We also encourage the use of whole-plasmid sequencing services such as Plasmidsaurus to verify integrity of the entire vector. [TROUBLESHOOTING] No colonies or high background, see **Supplementary Table 4**.

### Cloning custom transposon DNA (optional)

13) The section below provides optional cloning steps to truncate the VchCAST transposon right end to a 57-bp sequence (Option A) or replace the default payload sequence with a custom, user-desired payload (Option B). These steps can be performed before crRNA spacer cloning (steps 2-12) if needed, depending on the user’s application (e.g. inserting the same custom transposon at multiple different target sites).
  A. Truncating VchCAST transposon right end (57-bp) - [Timing - 2 d]
    i. Digest pDonor or pSPIN in the following reaction mix:

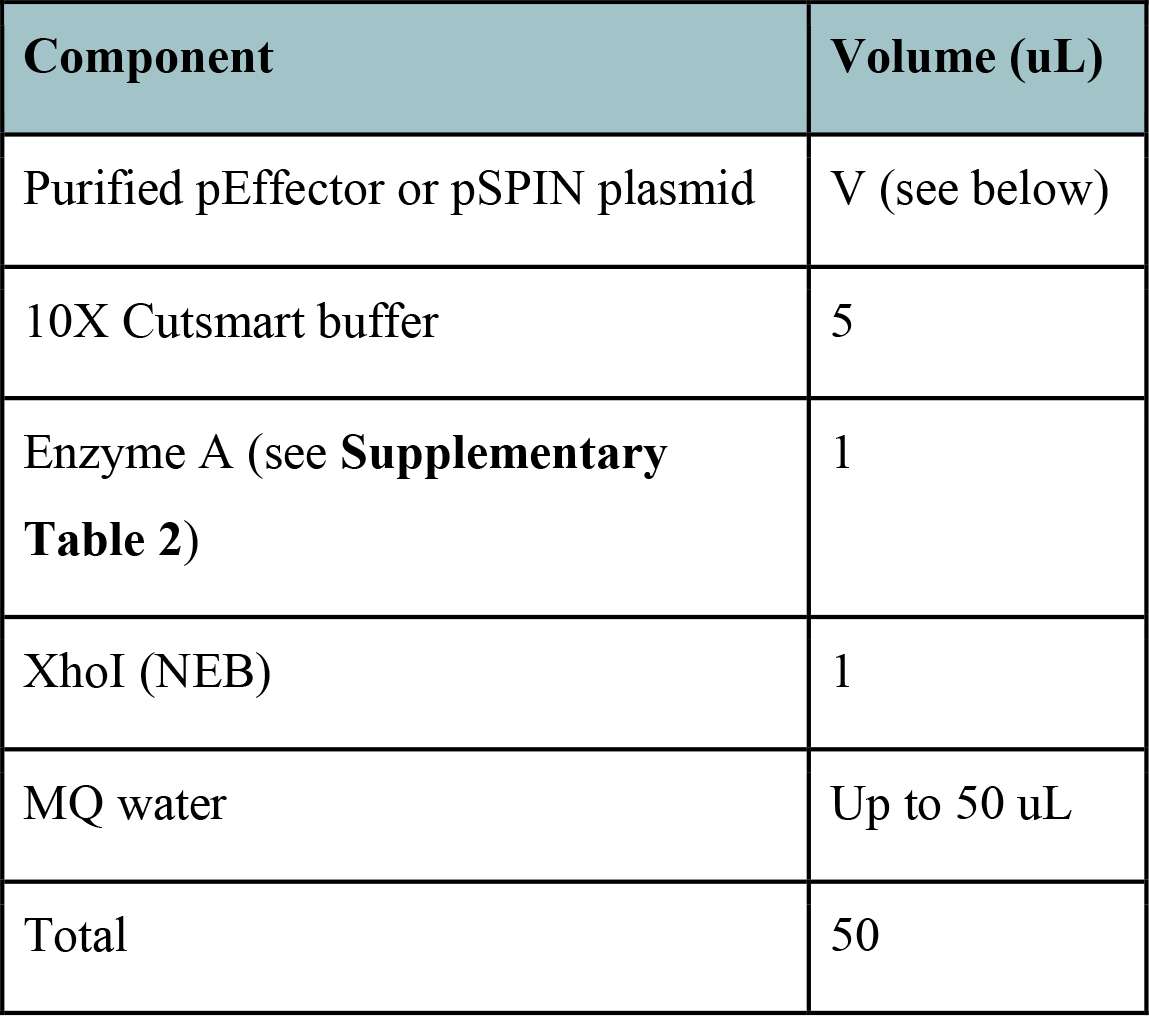 Digestion enzymes and suggested input DNA amounts:

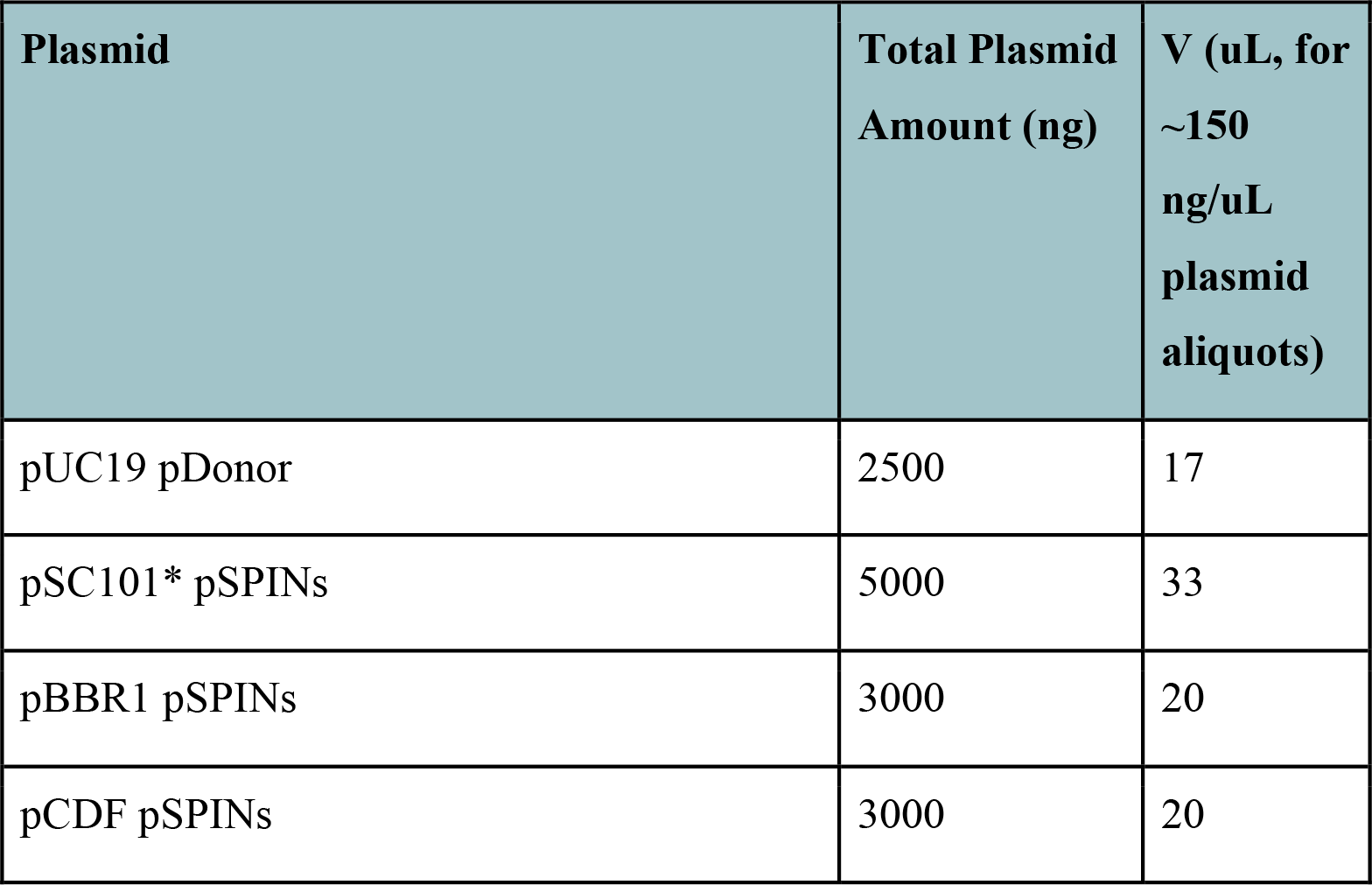 Incubate at 37 °C for 2 h.
    ii. Perform gel electrophoresis and extraction as described in Step 4-5. Expected band sizes are ∼12-14 kbp for pSPINs, and ∼3.3 kbp for pDonor.
    iii. Generate phosphorylated hybridized oligoduplexes separately for primer pairs (VchCAST_RE_fw1 / VchCAST_RE_rv1) and (VchCAST_RE_fw2/ VchCAST _RE_rv2) (**Supplementary Table 3**) as described in Steps 5-6.
    iv. Dilute each oligo duplex 1:50, and prepare ligation reactions as below. [CRITICAL] Prepare the ligation reaction on ice, or add the ligase last to prevent high levels of spurious ligation of the digested vector.

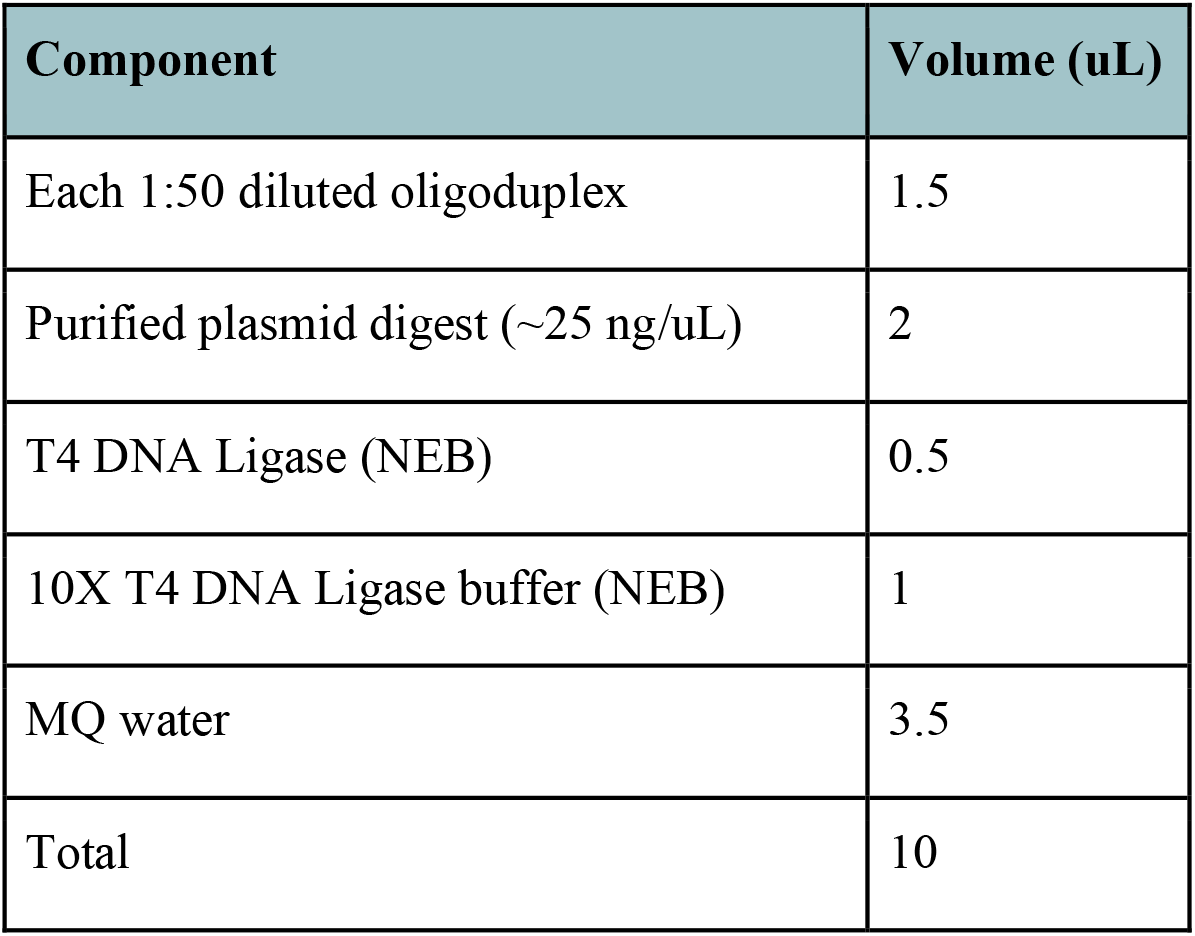 Incubate at room temperature for 30 min.
    v. Transform cells with entire ligation reaction, recover, plate, and inoculate colonies as in Steps 8-12. [TROUBLESHOOTING] No colonies or high background see **Supplementary Table 4**.
    vi. Confirm transposon right end sequence by Sanger sequencing (suggested primer is in **Supplementary Table 3**).
  B. Cloning custom transposon payload sequence - [Timing - 2 d]
    i. Digest pDonor or pSPIN in the following reaction mix:

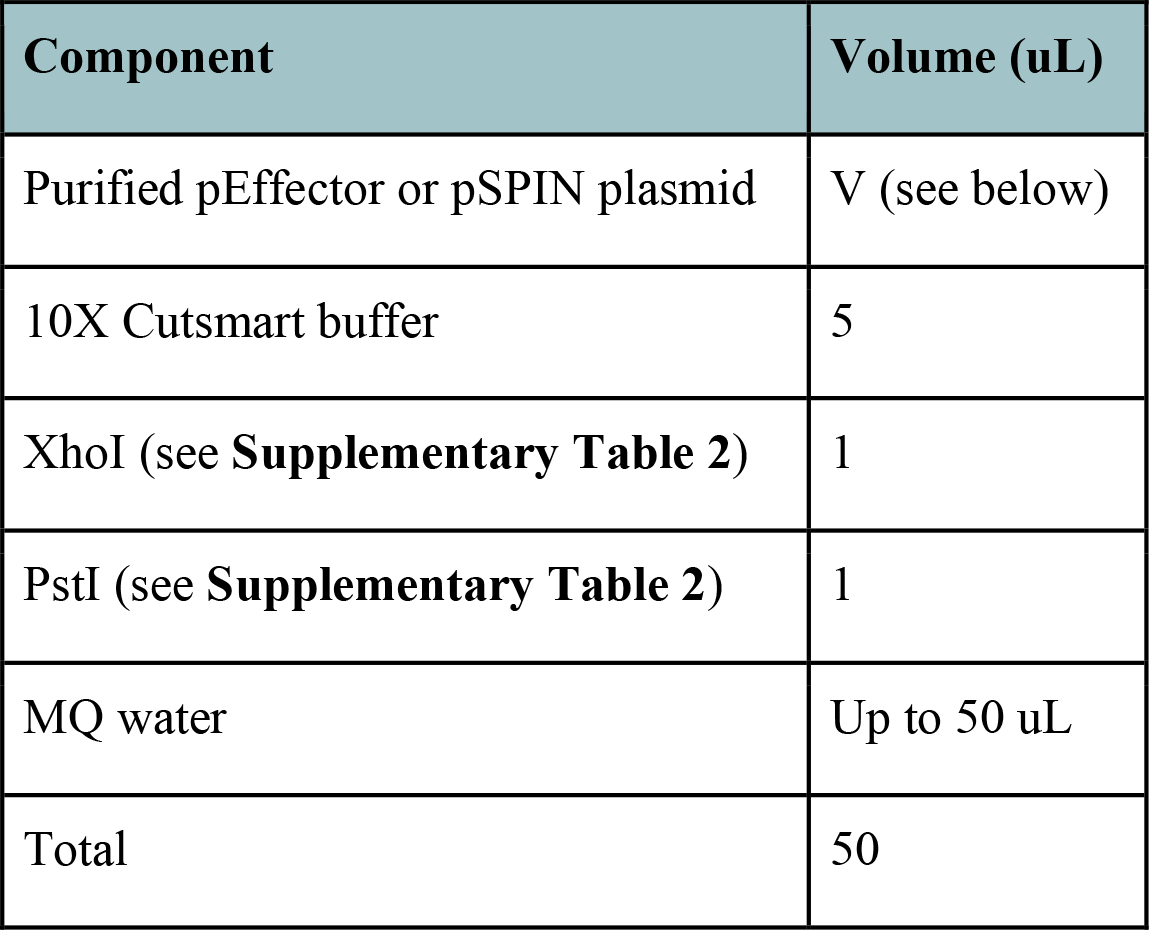 Suggested input DNA amounts:

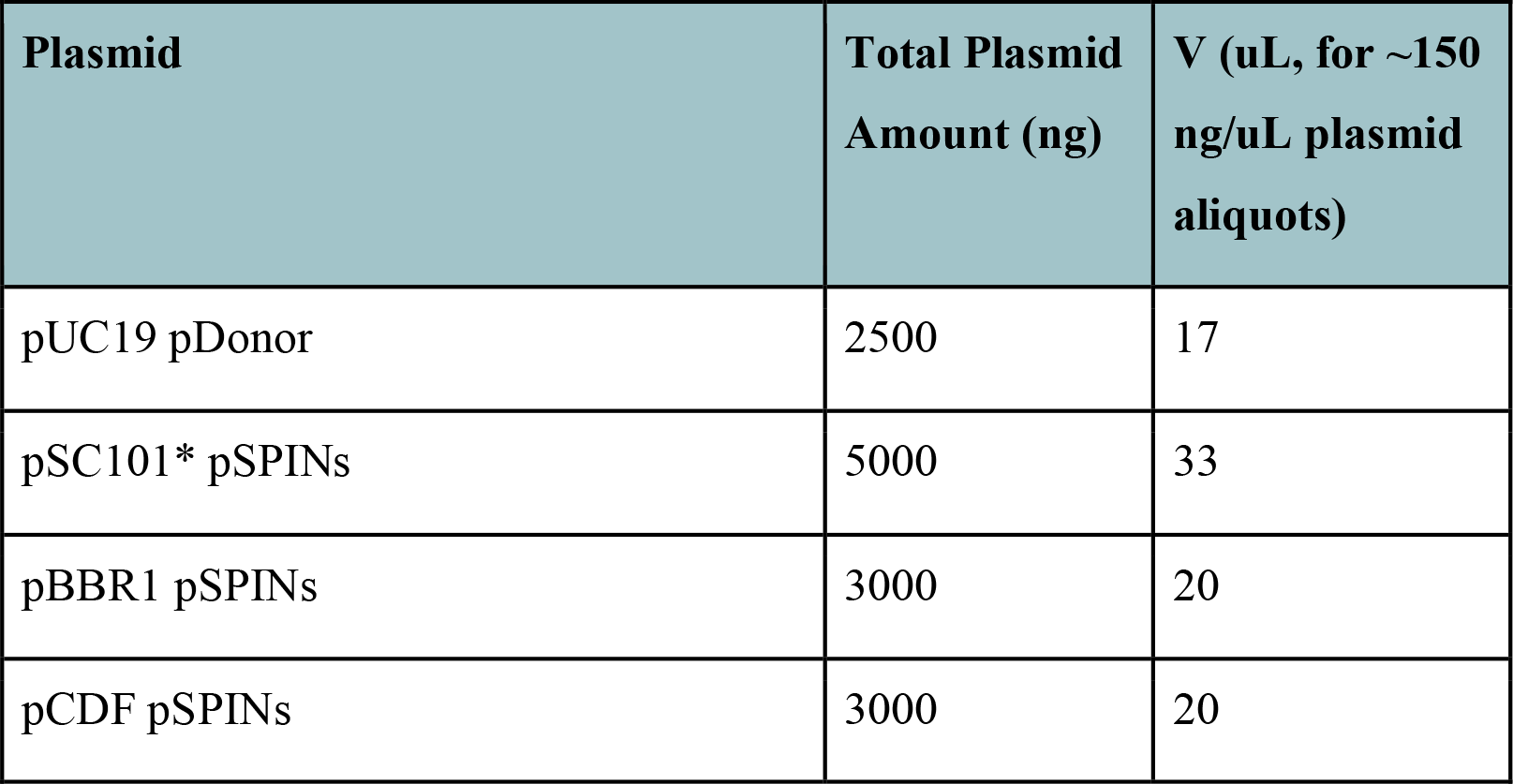 Incubate at 37 °C for 2 h.
    ii. Perform gel electrophoresis and extraction as described in Step 4-5. Expected band sizes are ∼11-13 kbp for pSPINs, and ∼2.7 kbp for pDonor.
    iii. Amplify custom payload DNA by PCR for Gibson cloning. Reaction mix below is customized for Q5 Hot Start Polymerase (NEB); a Q5 Hot Start 2X Mastermix, or similar high-fidelity polymerases or mastermixes can also be used instead by following manufacturer’s guidelines. [CRITICAL] Plasmid-derived template DNA should be diluted to <10 ng/uL; for amplifying from genomic DNA (gDNA) of bacteria, users should try a range of dilutions until a clean amplification is achieved. If amplifying from *E. coli* gDNA, users can adapt Steps 20Aiii in the analysis sections below.

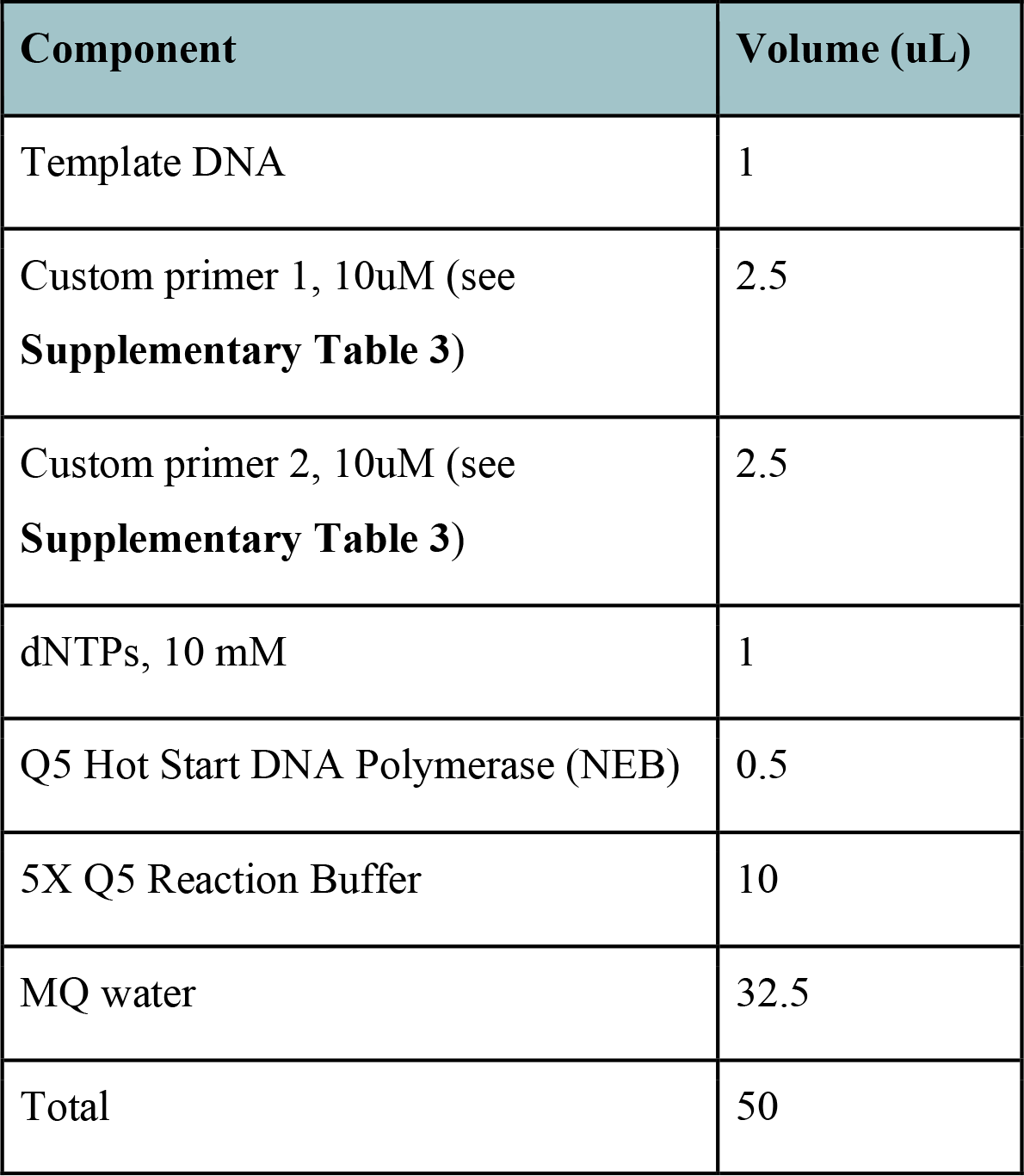
    iv. Perform PCR in a thermocycler with the following conditions:

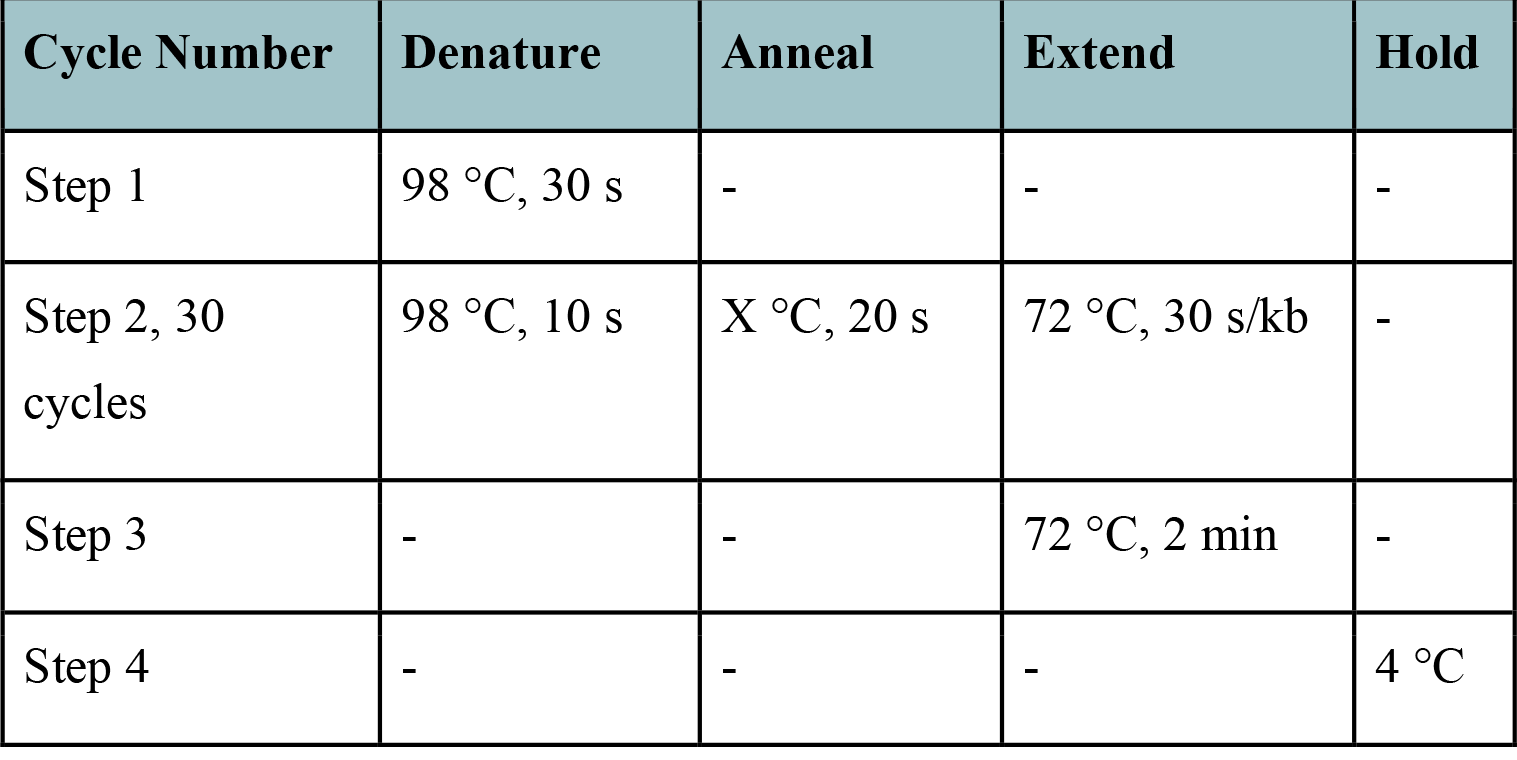 [CRITICAL] Vary temperature of annealing step according to primer binding sites, and if a different polymerase is used. Annealing temperature estimates can be found online at tmcalculator.neb.com.
    v. As described in Step 4-5, perform gel electrophoresis to confirm amplification of desired PCR product, excise correct bands and extract DNA. [PAUSE POINT]
    vi. In a PCR strip tube, prepare the Gibson assembly reaction below. [CRITICAL] For large multi-kb PCR inserts, increase the amount of purified PCR product in Gibson reaction.

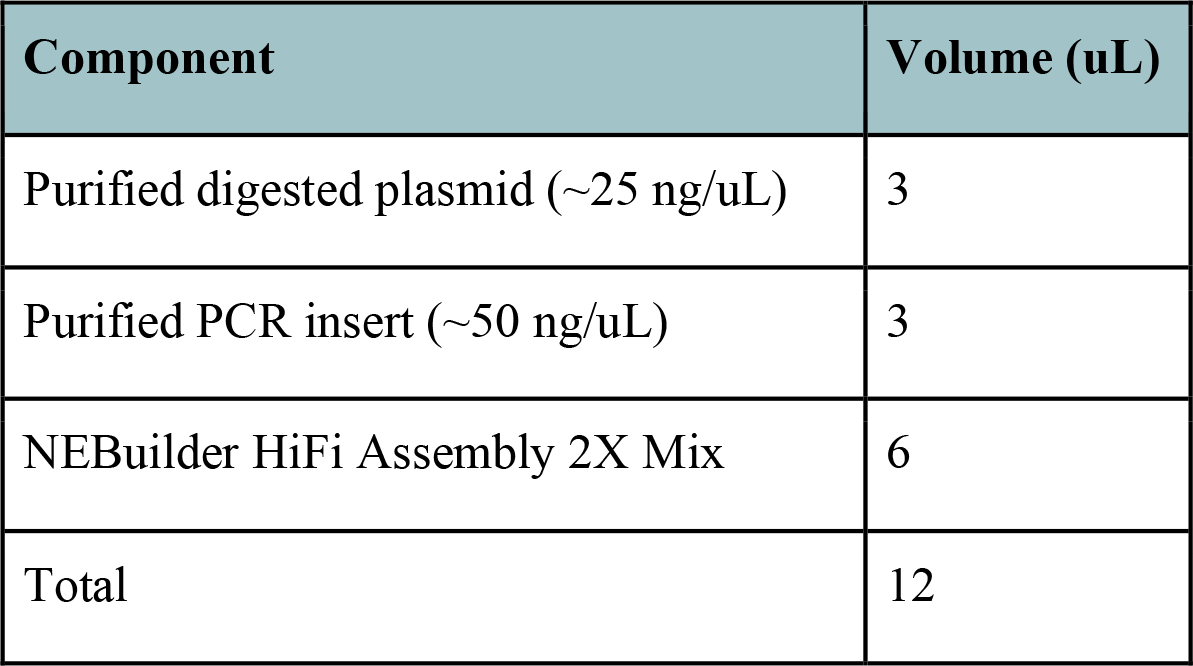 Incubate at 50 °C for 1-2 h in a thermocycler.
    vii. Transform cells with entire Gibson reaction, recover, plate, and inoculate colonies as in Steps 8-12. [TROUBLESHOOTING] No colonies or high background see **Supplementary Table 4**.
    viii. Confirm transposon payload sequence by plasmid sequencing (i.e., Plasmidsaurus).

### Perform transposition in target strain - [Timing - 3-4 d]

#### Transformation

14) pSPIN constructs can be delivered into cells using either chemical transformation (Option A) if targeting common *E. coli* strains, or electroporation (Option B) if targeting other strains or species. If using a pEffector+pDonor combination, electroporation is recommended in order to achieve sufficient transformation efficiency; however, chemical transformation can also be used, especially when using highly competent commercial *E. coli* cells. Alternatively, cells can be transformed first with either pEffector or pDonor, followed by inducing chemical transformation as in Step 2A, and then a single transformation with the second plasmid. As a negative control, we encourage users to transform in parallel using the respective non-targeting (NT) entry version of the vector constructs (prior to custom crRNA spacer cloning). This is an important control to test for non-CAST mediated integration and it should be expected to observe no integration at the target site.
  A. Deliver DNA by chemical transformation - [Timing - 2-3 d]
    i. On ice, mix ∼200-300 ng of each construct plasmid with 50 uL of chemically competent cells (commercial Sigma BL21 CMC0014 or homemade for other bacteria). If performing a two-plasmid transformation, use 300-400 ng of each plasmid.
    ii. Incubate on ice for 15 min, heat shock at 42 °C for 30s, then return to ice for 5 min.
    iii. Add each transformation to a microcentrifuge tube with 950 uL LB media. Recover with shaking or rotation at 37 °C for 1-2 h. [CRITICAL] Recover at 30 °C for 2 h if using pSC101* constructs.
    iv. Plate 100 uL of recovery on an LB agar plate containing appropriate antibiotic selection (**Supplementary Table 1**). Pellet the remaining 900 uL recovery, discard supernatant, resuspend in 100 uL LB and plate on a similar LB agar plate. Ensure that all visible liquid pools have dried off from plates, and incubate plates at 37 °C overnight for 18-24h. [CRITICAL] Plates can alternatively be incubated at 30 °C for 28-30 h, or at 25 °C for 40-48 h, which may induce higher integration efficiencies. Incubate plates at 30 °C or lower if using pSC101* constructs.
  B. Delivery by electroporation - [Timing - 2-3 d]
    i. Pre-chill 1 mm electroporation cuvettes on ice. Pre-warm 950 uL of LB media in microcentrifuge tubes at 37 °C using a heat block or incubator.
    ii. On ice, mix ∼50-100 ng of each construct plasmid with 40 uL of electro competent cells (commercial Sigma BL21 CMC0016 or homemade for other bacteria). If performing a two-plasmid transformation, use 100-200 ng of each construct plasmid.
    iii. Pipette cell and plasmid mixture into a cold electroporation cuvette. Electroporate with a GenePulser electroporator (Bio-Rad) or equivalent at 2000 kV, 25 uF and 200 Ohms. Arc times in the range of 4.5 – 5.1 are considered a highly efficient electroporation.
    iv. [CRITICAL] Immediately pipette 700 uL of pre-warmed LB media from a microcentrifuge tube, mix by pipetting and transfer entire mixture back into the tube. Delays in adding recovery media to electroporated cells can reduce transformation efficiencies.
    v. Recover cells with shaking or rotation at 37 °C [CRITICAL] Recover at 30 °C for 2 h if using pSC101* constructs.
    vi. Plate cells on LB agar and grow overnight at 37 °C as in Step 15iv. [CRITICAL] Plates can alternatively be incubated at 30 °C for 28-30 h, or at 25 °C for 38-44 h, which may induce higher integration efficiencies. Incubate plates at 30 °C or lower if using pSC101* constructs.

#### Conjugation

15) A conjugative method for delivery is only viable with pSPIN constructs that are engineered with an origin of transfer, a necessary component for conjugation, and delivered with a conjugative strain that harbors genetically encoded RP4 conjugative machinery. This method is especially useful for delivery into strains that are recalcitrant to transformation. The delivery method is reliant on a donor strain transformed with pSPIN. A commonly used donor strain is *E. coli* EcGT2, which utilizes RK2-based conjugal transfer^82^. This *EcGT2* donor strain is auxotrophic for the essential cell-wall component diaminopimelic acid (DAP), thus requiring DAP supplementation (50 ug/mL) in the growth media. This allows for counterselection of the donor after conjugation. As a negative control, transform in parallel using the respective NT entry version of constructs (before custom crRNA spacer cloning).
  A. Generate chemically competent EcGT2 donor strain harboring pSPIN chemical transformation– [Timing – 2d]
    i. Inoculate 10 mL LB media with EcGT2 from colonies or a frozen glycerol stock, and grow with shaking at 37 °C overnight. Supplement media with 1X concentration of the appropriate antibiotics if required for the target strain.
    ii. The following day, make a 1:100 dilution of the overnight culture into 100 mL of LB media in a 250 mL sterile baffled Erlenmeyer flask. Supplement media with antibiotics if needed. Each 100 mL culture produces enough competent cells for ∼80 50-uL transformation reactions. If larger preparations are required, the volume can be scaled up (to 1 L of media) using a 2 L flask.
    iii. Incubate with shaking at 37 °C. Measure OD600 of the culture regularly
    iv. Once cells reach an OD600 of ∼0.5 (approximately 1–2.5 hrs), transfer the culture into 50 mL conical tubes with 50 mL of culture per tube, and incubate tubes on ice for 15 min. [CRITICAL] The following steps should be performed with cells on ice as much as possible. All centrifugation steps below should be performed at 4 °C.
    v. Pellet cells at 4000 g for 10 min and discard supernatant.
    vi. Add 20 mL of ice-cold Buffer A into each conical tube, and resuspend pellet fully by pipetting up and down. Repeat centrifugation step v)
    vii. Add 2 mL of ice-cold Buffer B into each conical tube, resuspend fully by pipetting. At this step, cell resuspensions from multiple tubes can be combined into one tube.
    viii. Add 70 uL DMSO per 2 mL of resuspension. Mix gently and incubate on ice for 15 min. Put PCR strip tubes or microcentrifuge tubes on ice.
    ix. After 15 min, add an additional 70 uL of DMSO per 2 mL of resuspension and mix gently.
    x. Aliquot 100 uL of cell resuspension per cold PCR strip tube, or larger volumes into cold microcentrifuge tubes.
    xi. Transform cells immediately (Step xiii below), or flash freeze in liquid nitrogen and store at -80 °C for up to 12 months. Freezing cells usually leads to decreased transformation efficiencies.
    xii. On ice, mix 20-100 ng of each CAST plasmid construct (pSPIN) with 50 uL of chemically competent EcGT2 cells.
    xiii. Incubate on ice for 15 min, heat shock at 42 °C for 30s, then return to ice for 5 min.
    xiv. Add each transformation to a microcentrifuge tube with 950 uL LB media. Recover with shaking or rotation at 37 °C for 1 h.
    xv. Plate 100 uL of recovery on an LB agar plate containing the appropriate antibiotic and DAP (**Supplementary Table 1**). Pellet the remaining 900 uL recovery, discard supernatant, resuspend in 100 uL LB and plate on a similar LB agar plate. Incubate plates at 37 °C overnight for 12-18h.
    xvi. The next day, pick a few colonies of EcGT2 pSPIN transformant in LB liquid media supplemented with appropriate antibiotic and DAP and grow overnight.
    xvii. Do the same as step xvi for the recipient strain but do not supplement DAP. Now that the donor strain harboring a CAST plasmid of interest has been generated, delivery of pSPIN from the donor into the recipient strain for subsequent editing can be carried out in the next step.
  B. Conjugative delivery of pSPIN – [Timing – 1 d]
    i. Pipet 1.5 mL of EcGT2 donor strain into a 1.7 mL microcentrifuge tube, and do the same for recipient strain in a separate tube.
    ii. Spin down aliquoted cultures in a centrifuge for 5 minutes at 4,000 x g at RT.
    iii. Decant supernatant and aliquot another 1.5 mL of donor or recipient on top of the remaining pellet in each respective tube. Resuspend the pellet.
    iv. Spin down aliquoted cultures in a centrifuge for 5 minutes at 4,000 x g at RT.
    v. Decant supernatant and add 1 mL of 1x PBS pH 7.4 to each tube. Resuspend pellets to wash them.
    vi. Spin down aliquoted cultures in a centrifuge for 5 minutes at 4,000 x g at RT.
    vii. Decant supernatant and add 1 mL of 1x PBS pH 7.4 to each tube.
    viii. Repeat this wash for a total of 3x washes.
    ix. After the final wash, resuspend donor and recipient pellets in 1 mL of 1x PBS pH 7.4.
    x. Calculate number of cells either by OD_600_ or flow cytometry-based count.
    xi. Aim to have ∼ 8x10^8^ – 1x10^9^ cells for both donor (*E. coli* OD_600_ = 1) and recipient, to have a final 1:1 ratio of donor:recipient. [CRITICAL] If the recipient is recalcitrant to conjugation, it is recommended to increase the donor:recipient ratio to 10:1, in order to increase conjugation efficiency.
    xii. In a new 1.7 mL microcentrifuge tube, add the required volumes corresponding to the adequate amount of cells for both donor and recipient, combining them together. This is your conjugation reaction. If conjugation volumes exceed >1.7 mL of culture solution, further concentrate your cells by centrifugation (Step vi) and resuspend in a smaller volume of PBS.
    xiii. As controls, add the same volumes to individual tubes containing only the donor or only the recipient with no additional solutions added.
    xiv. Spin all tubes down in a centrifuge for 5 minutes at 4,000 x g at RT.
    xv. Aspirate all supernatant carefully, leaving behind only a cell pellet.
    xvi. Carefully resuspend pellets in 10 uL of 1x PBS pH 7.4. [CRITICAL] Be sure not to introduce any bubbles in the mixture, as the solution will be highly viscous.
    xvii. Pipet the 10 uL cell mixture as a uniform spot onto LB plates supplemented with DAP. [CRITICAL] Do not introduce any bubbles in the spot as this will disrupt conjugation (cell-to-cell contact). In addition, different media conditions (non-LB) should be considered here, depending on the recipient strain.
    xviii. Allow spots to dry for up to 30 min.
    xix. Incubate plates at 37 °C for 16–24 h. [CRITICAL] longer time for conjugation can increase the conjugation efficiency.
  C. Selection of transconjugants – [Timing – 1-2 d]
    i. After 16–24 h of growth, spot reactions will have fully grown.
    ii. Scrape the spot reactions with an inoculation loop and mix into 1 mL 1x PBS pH 7.4 in a 1.7 mL microcentrifuge tube.
    iii. Resuspend reaction fully with a P1000 pipet.
    iv. Make serial dilutions from 10^0^ – 10^-7^ in a 96-well plate, in which the dilutions comprise 20 uL of sample combined with 180 uL of 1 mL 1x PBS pH 7.4. The dilutions are described in the Table below. Make dilutions for each conjugation samples, and for the donor only and recipient only samples.

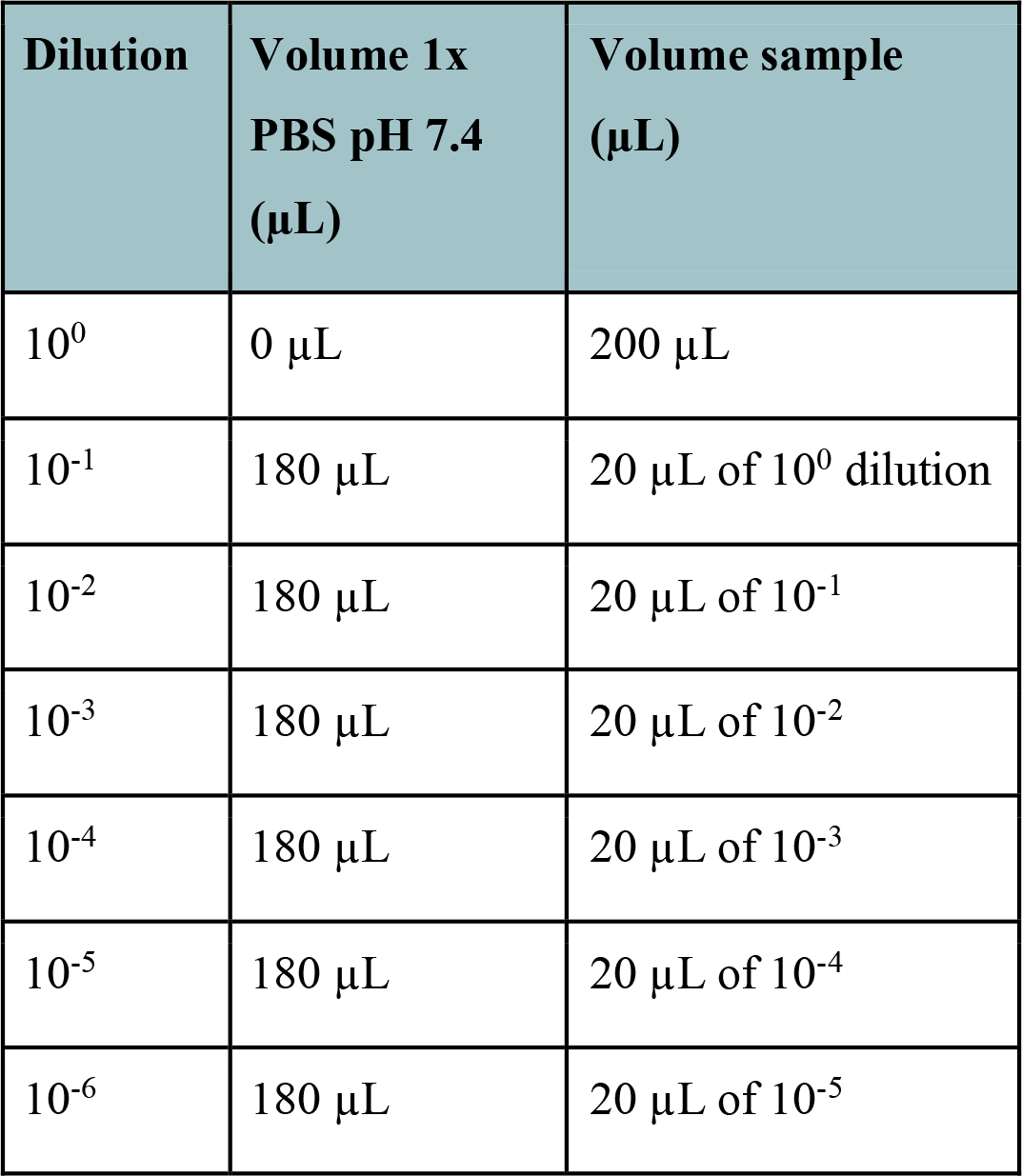
    v. Thoroughly mix dilutions using a p200 multi-channel pipet.
    vi. Plate 5 uL spots of dilutions for each sample using an 8-well multi-channel pipet onto:
      1. LB plates without antibiotics,
      2. LB supplemented with the appropriate antibiotic encoded on the plasmid backbone (i.e. Kan),
    vii. [CRITICAL] Do not supplement any plates for selection of transconjugants with DAP because inclusion of DAP will allow for the growth of the donor strain and convolute downstream results downstream.
    viii. Let spots dry for up to 30 min.
    ix. In addition, plate 100 uL of resuspended conjugation reactions on an LB agar plate containing appropriate antibiotic selection. These plates will be used for acquiring biomass to estimate integration efficiency.
    x. Incubate all plates at 37 °C for 12–24 h.
  D. Alternative selection of transconjugants if conjugation efficiency is already known for target bacteria:
    i. After 16-24 h of growth, spot reactions will have fully grown.
    ii. Scrape the spot reactions with an inoculation loop and mix into 1 mL 1x PBS pH 7.4 in a 1.5 mL microcentrifuge tube.
    iii. Resuspend reaction fully with a P1000 pipet.
    iv. Make serial dilutions 10^0^ – 10^-4^/10^-6^ depending on the expected conjugation efficiency.
    v. Plate 100 uL of each dilution
    vi. [CRITICAL] if very low efficiency is expected, spin down the conjugation 5 minutes at 4,000 x g at RT, resuspend in 100uL and plate all.
    vii. [CRITICAL] Do not supplement any plates for selection of transconjugants with DAP because inclusion of DAP will allow for the growth of the donor strain and convolute results downstream.
    viii. Incubate all plates at 37 C for 12-72 h depending on the growth rate of the recipient strain.

### Extract genomic DNA from transformants/transconjugants in bulk and passage for clonal isolation - [Timing - 3-4 d]

We recommend assessing integration efficiency across the entire population of cells as a quality control check for CAST activity. In addition, colonies at this stage will typically not be clonal after a single transformation step, and thus requires passaging repeatedly if the goal is clonal isolation.

16) To conduct bulk gDNA extraction, after incubation on LB agar, select one of the two plates from each transformation that produced >100 colonies, but not a dense lawn of cells. Scrape at least 100 colonies from the plate using a pipette tip or inoculation loop, and fully resuspend in 500 uL of LB media. [CRITICAL] If very few colonies are produced from step 15, they can be re-streaked onto a new agar plate and incubated overnight to produce enough cell material for Step 16. If both plates produced dense lawns of cells where individual colonies are not easily discernible, plate lower dilutions of the recovery. Dense lawns may reduce integration efficiencies. [TROUBLESHOOTING] Few or no colonies on transposition plates see **Supplementary Table 4**.
17) Prepare 1:10 serial dilutions of cell resuspension into 100 uL aliquots of LB media and plate each dilution onto an LB agar plate with appropriate selection and incubate overnight. This is required for clonal isolation (see Experimental Design: V. DNA integration analysis). For clonal isolation, go to step “Isolating clonal integrants by PCR”.
18) (Optional) Repeat step 16 and 17 once or several times, which may help induce higher integration efficiencies.
19) Genomic DNA extraction for PCR and qPCR.
  i. Add 100 uL of resuspended cells from step 17 or 18 into 900 uL of LB media in a plasmid cuvette, and measure OD_600_. Dilute cell suspension or scrape and resuspend more colonies until the OD_600_ reaches ∼0.4-0.5. [CRITICAL] The cell suspension has roughly equivalent cell material as a turbid overnight culture and can be used for gDNA extraction using the Wizard gDNA Purification kit (Promega) or equivalent. However, for general PCR/qPCR, heat lysis as described below is also sufficient to produce template DNA.
  ii. Pipette 170 uL of each cell resuspension into a microcentrifuge tube. If many parallel resuspensions are involved, PCR strip tubes or PCR plates can be used instead, with 85 uL of resuspension per tube or well.
  iii. Pellet at 12,000 x g for 2 min for microcentrifuge tubes, or 4000 g for 5 min for PCR strips or PCR plates.
  iv. Discard supernatant and fully resuspend pellet in 160 uL of MQ water for microcentrifuge tubes, or 80 uL for PCR strips or PCR plates.
  v. Heat resuspensions at 95 °C for 10 min using a heat block or thermocycler. For microcentrifuge tubes, seal caps with parafilm or plastic cap lock.
  vi. Cool to room temperature and pellet as in step iii. For each microcentrifuge tube, dilute 10 uL of supernatants into 390 uL of MQ water in a new microcentrifuge tube; for PCR strip tubes, dilute 2 uL into 78 uL of MQ water into new PCR strips or PCR plates. [PAUSE POINT] Diluted lysates can be stored for several months at -20 °C.

### (Optional) Population-wide PCR and qPCR analysis of transposon integration - [Timing - 3 h]

20) Genomic DNA extracts prepared in step 19 can be used for population-wide PCR analysis to confirm successful integration (option A), or for measurement of integration efficiencies by qPCR (option B). These steps may be useful in troubleshooting or optimizing experimental conditions for transposition, especially for new target sites or new target hosts, before proceeding to clonal integrant selection.
  A. Population PCR analysis. [CRITICAL] We recommend performing three PCR reactions for target site and sample: one pair each probing for the two possible insertion orientation (T-RL or T-LR), and a third control pair probing for an unmodified (wild-type) genomic locus.
    i. Follow **Fig. 5A-B** and **Supplementary Table 3** for junction PCR primer design.
    ii. [CRITICAL] For genotyping, high-fidelity polymerases such as Q5 (NEB) are recommended. However, lower-fidelity polymerases such as OneTaq (NEB) or the equivalent can also be used. [CRITICAL] If only integration in the T-RL is desired, there is no need to detect integration in the T-LR orientation. However, PCR to detect T-LR can be performed in parallel using a new pair of primers specific to the T-LR junction (**Supplementary Table 3**). Set up a PCR reaction for gDNA extracts as below. Preparation of a batch mastermix is recommended for multiple samples in parallel.

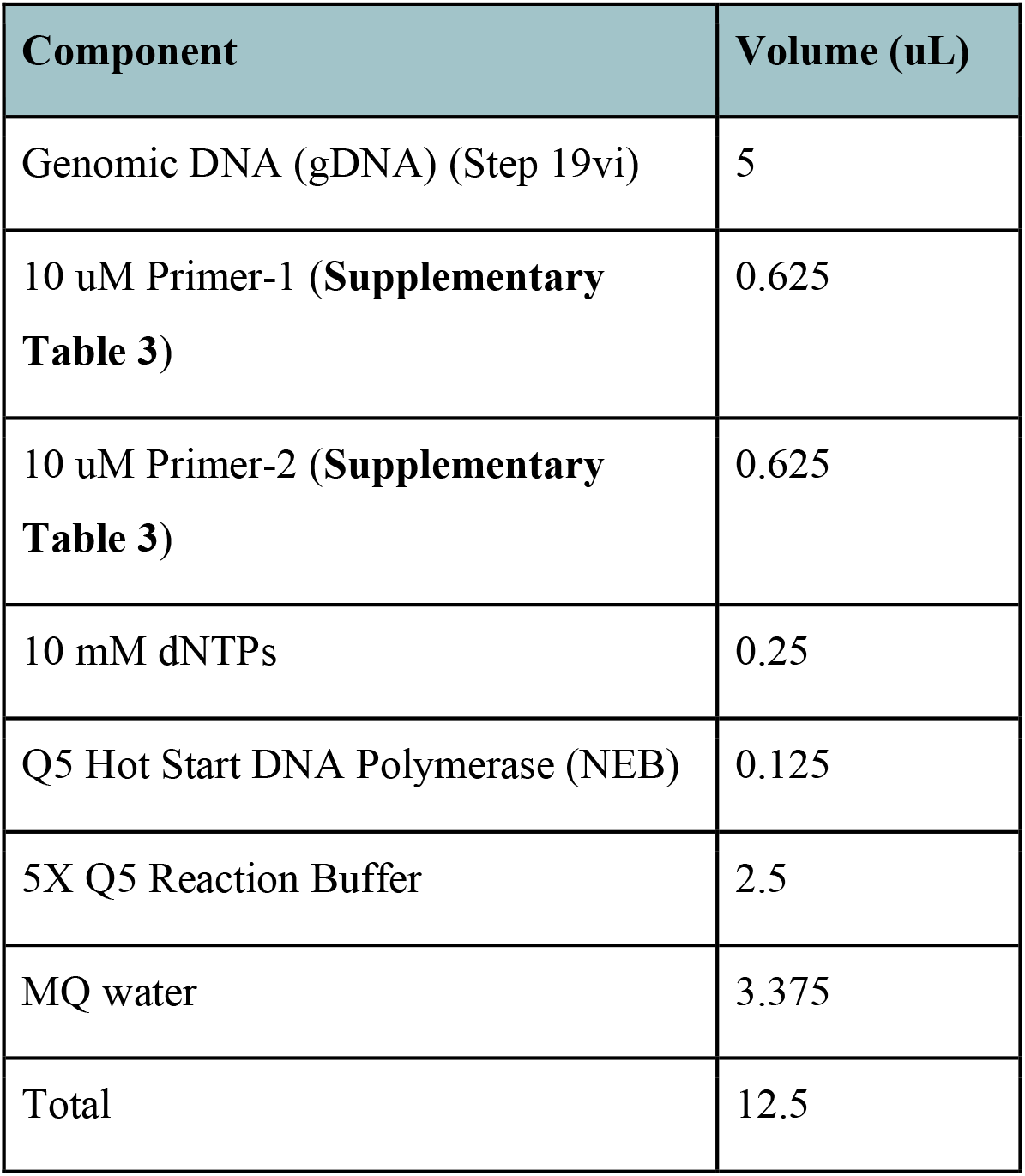
    iii. Perform PCR in a thermocycler with the following conditions:

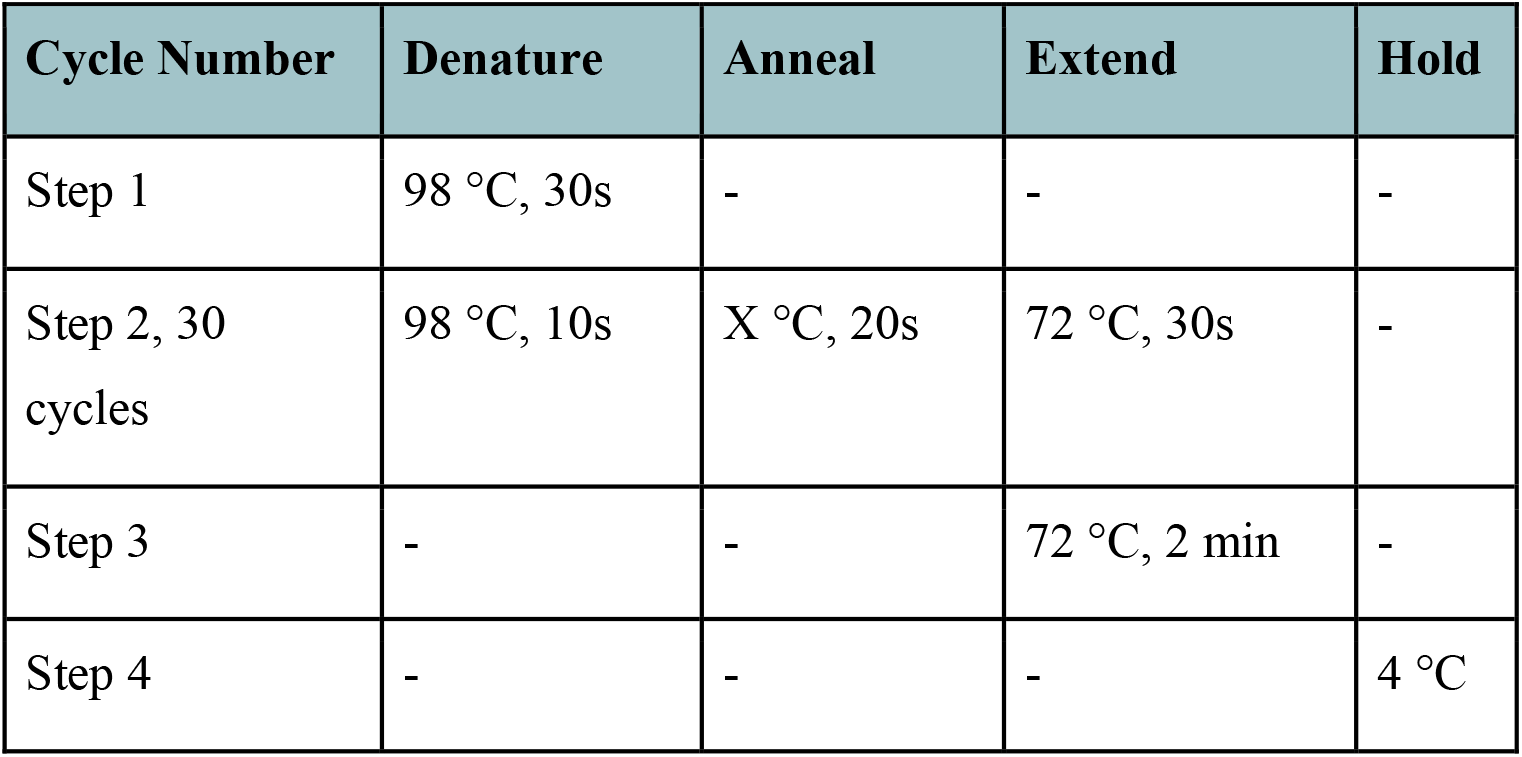 [CRITICAL] Vary temperature of annealing step according to primer binding sites and polymerase used. Annealing temperature estimates can be found online at tmcalculator.neb.com. Increase extension time for longer expected PCR products (for Q5 Polymerase, ∼30s of increased extension is recommended for each additional 1 kb of product).
    iv. Perform gel electrophoresis as described in Step 4. For PCR products smaller than 1 kb, a 1.5% agarose gel is recommended. Confirm the existence of a band at the correct size corresponding to a transposon-genome junction product. [TROUBLESHOOTING] No PCR/qPCR bands detected, see Supplementary Table 4.
    v. (Optional) If further confirmation is necessary of a bona fide integration event, it is recommended to gel extract and sequence the target band. Using a clean blade or razor, cut out the desired band and gel extract using the QIAquick or MinElute gel extraction kits (Qiagen).
    vi. (Optional) Confirm insertion by Sanger sequencing of extracted bands; either PCR primer (from step 20Ai) can be used for sequencing.
  B. Population qPCR analysis.
    i. Follow **Fig. 5A-B** and **Supplementary Table 3** for qPCR primer design considerations. We typically recommend qPCR primers of 18–24 nt in length, with an average predicted melting temperature of 55 °C. [CRITICAL] The amplification efficiency of each qPCR primer pair should be determined before use. This can be done by measuring Cq values generated from a serial dilution series of the sample lysate or gDNA. Plots of Cq values vs log dilution should produce a straight line with a negative slope between -3.10 and -3.60.
    ii. Prepare diluted primer mixture by adding 10 uL of primer 1 and 10 uL of primer 2 (**Supplementary Table 3**) to 380 uL of EB buffer or MQ water.
    iii. Set up a qPCR reaction for each diluted lysate as below in a 384-well qPCR plate. Preparation of a batch mastermix is recommended for multiple samples in parallel.

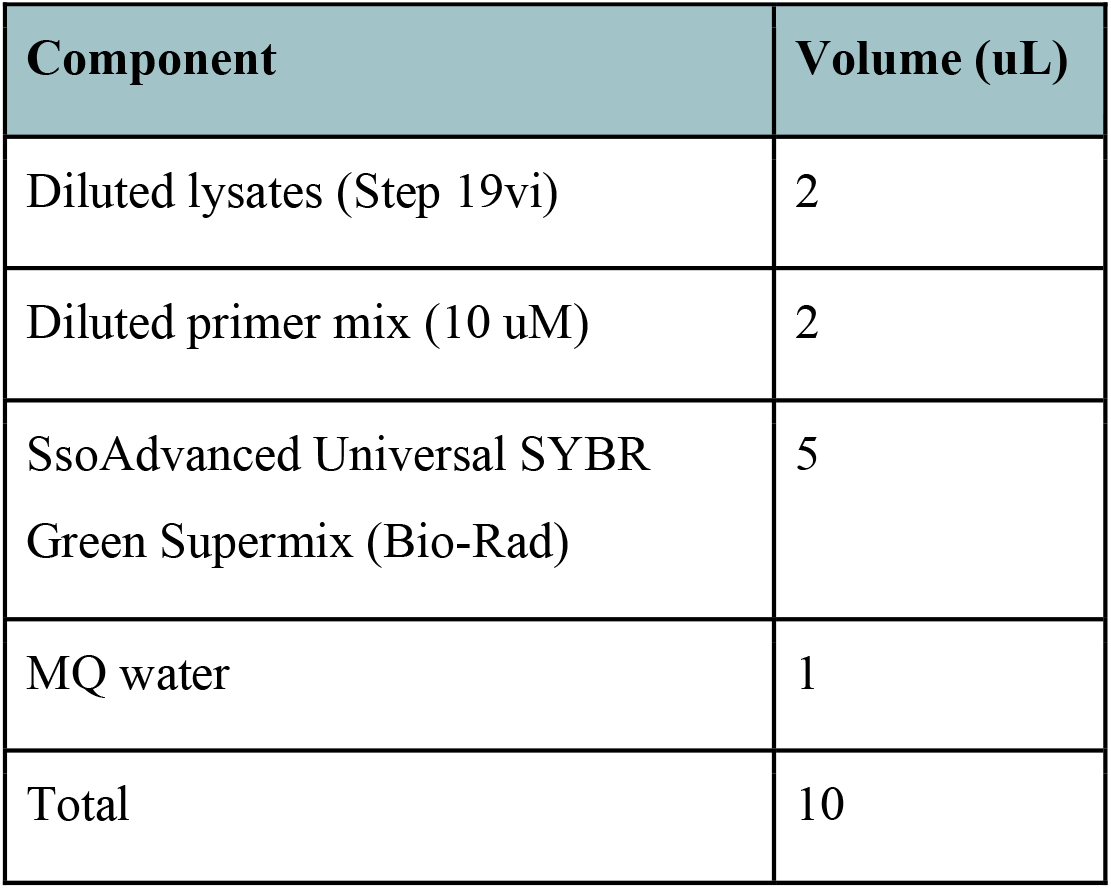 [CRITICAL] For each sample, perform qPCR with primer pair RL and pair G (**Supplementary Table 3**). If integration efficiency in the T-LR orientation is needed, perform a third parallel qPCR reaction with primer pair L. (Optional) Perform each reaction in three separate technical replicates. We strongly recommend including crRNA-NT and water-only samples as negative controls.
    iv. Perform qPCR in a 384-well qPCR thermocycler with the following conditions:

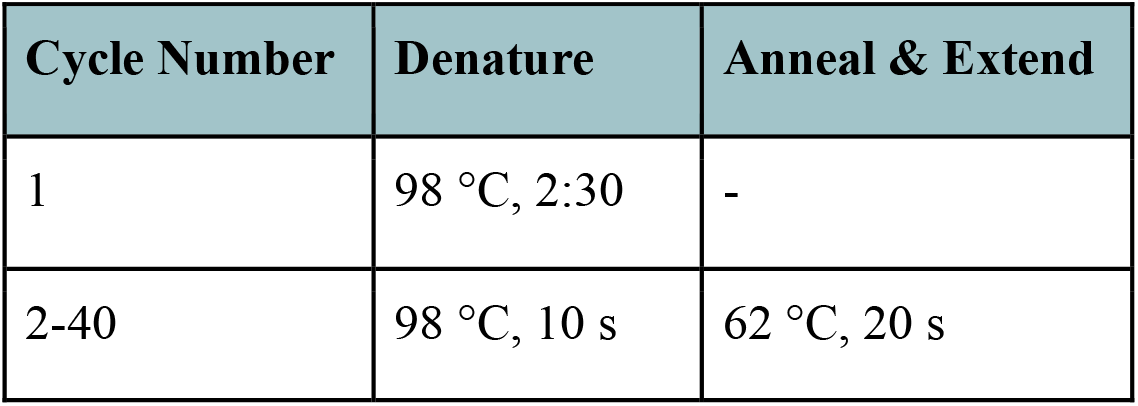 [CRITICAL] Vary the annealing and extension for new primer pairs as needed to obtain sufficiently high amplification efficiencies, without background off-target amplifications.
    v. Calculate estimated transposition efficiency as in **Fig. 5A-B** and **Box 3**. [TROUBLESHOOTING] Low transposition efficiencies, see **Supplementary Table 4**.
    vi. (Optional) Mix each entire reaction with 6X loading dye, load and perform gel electrophoresis using a 1.5% agarose gel. Typical electrophoresis conditions for a ∼200 bp product are 130 V for 22 min. [TROUBLESHOOTING] No PCR/qPCR bands detected, see **Supplementary Table 4**.
    vii. (Optional) Excise bands at the expected sizes for reactions RL and LR, and extract DNA as in Step 4. An elution of 10 µL buffer EB using the MinElute Extraction Kit (Qiagen) is recommended. Confirm sequence by Sanger sequencing using one of the primers in the primer pair.

#### Box 3. qPCR-based calculation of DNA integration efficiencies

Integration efficiency measurements can be made for a given insertion orientation (e.g. T-RL), using Cq values from a primer pair ‘RL’ probing for the T-RL orientation, and a primer pair ‘G’ probing for a reference genomic locus, as follows:

%T-RL = 100 x 2^^Cq(Pair G) - Cq(Pair RL)^

Similarly, the % integration in the T-LR orientation can be calculated using the Cq value for the T-LR primer pair. The total integration efficiency at the target site is then calculated as the sum of integration efficiencies in both of the two possible orientations. We note that, using this method, the error magnitiude (PCR amplification efficiencies between different loci) increases as efficiencies approach 100%, and it is possible to obtain artifactual values larger than 100%. In such scenarios, we recommend testing multiple different primer pairs and/or repeating the qPCR with additional biological and technical replicates.

A more definite measurement of integration efficiency can be obtained by normalizing against a clonal integrant. Cells containing clonal insertions should first be isolated (Steps 23-31), which theoretically have an integration efficiency of 100% in one orientation. qPCR analyses are then performed as above on both the clonal strain and the unknown sample(s), followed by normalization. For example, the T-RL integration efficiency can be calculated as:

True %T-RL = 100 x (T-RL sample)/(T-RL clonal)

### (Optional) Estimation of conjugation efficiency - [Timing - 1 h]

21) Using serial dilutions of conjugation reactions in Step 15C, an estimation of conjugation efficiency from donor to recipient is possible by comparing colony counts between conjugation reactions, donor-only reactions, and recipient-only reactions.
  A. Colony counting of conjugation reactions
    i. After 12–24 h incubation of spots, remove plates from incubator.
    ii. Count the number of colonies for the two highest dilution series that grew on LB only and LB supplemented with the appropriate antibiotic encoded on the plasmid backbone (i.e. Kan), for conjugation reactions, donor-only reactions, and recipient-only reactions. For the number of colony forming units (CFU) in the selective plates (R) and the number of colony forming units (CFU) in non-selective plates (C), at 10^*n*^ dilutions with *n* = the fold dilution in the series of dilutions plated (i.e., 10^-3^), apply this calculation:

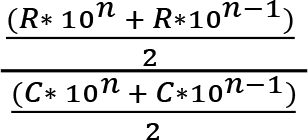
    iii. [CRITICAL] If LB plates were incubated without DAP supplementation, it is crucial that no growth is observed for donor-only reactions. If growth is observed, conjugation efficiency results will be inconclusive.
    iv. [CRITICAL] Growth is expected for transconjugants and for recipient-only reactions on LB plates, but only growth of transconjugants is expected on LB supplemented with the appropriate antibiotic. If recipient-only reactions exhibit growth on the antibiotic, it is necessary to determine the minimum inhibitory concentration (MIC) and repeat the experiment at that concentration of antibiotic.
22) If donor-only and recipient-only colony counts correspond to expected results in 21Aiii-iv, proceed with conjugation efficiency calculations. For the transconjugant, divide the number of colonies in the highest dilution on LB only plates by the number of colonies in the highest dilution on LB supplemented with appropriate antibiotic. This final number is the conjugation efficiency.

### Isolating clonal integrants by PCR - [Timing - 18 h]

The steps below describe isolation of clonal integrants from steps 17 or 18, achieved by genotyping random colonies using colony PCR. We want to emphasize that if the goal is clonal isolation, colonies will typically not be clonal after a single transformation step, and thus requires passaging repeatedly (at least once) to homogenously fix the integration product. Depending on genomic insertion site or transposon payload genes, users may instead opt to isolate clonal integrants through phenotyping, then subsequently confirm insertions by PCR. For detection of clonal insertions, an external-external PCR strategy is used (**Fig. 5C-D**). We recommend performing an additional PCR reaction for an unmodified genomic locus (as in Step 20A-B), as a positive control that should yield a band for the PCR reaction.

[CRITICAL] For multiplexed experiments, selection for clonal integration can be performed by performing genotyping PCR at all target sites simultaneously. If colonies with clonal insertion at all sites are not found from the first round of PCR, colonies with clonal insertions at several sites can be re-streaked for an additional overnight incubation, followed by further PCR genotyping.

23) From single colonies derived from plates after Steps 17 and 18, pick 10–20 colonies. Resuspend the cell material for each colony in 40 uL of MQ water. [CRITICAL] Include one or two colonies from a negative control plate.
24) Onto a new LB agar plate with appropriate antibiotic selection, spot 1 uL of each cell resuspension. Let the spots dry completely, then incubate plate at 37 °C overnight. [CRITICAL] Incubate plate at 30 °C if using pSC101* constructs.
25) Heat the cell resuspension at 95 °C for 10 min,and then cool to room temperature. Perform a 1:20 dilution of each lysate into 40 uL of MQ water.
26) [CRITICAL] For genotyping, high-fidelity polymerases such as Q5 (NEB) are recommended. However, lower-fidelity polymerases such as OneTaq (NEB) or equivalent can also be used. Set up a PCR reaction for each diluted lysate as below. Preparation of a batch mastermix is recommended. Follow **Fig. 5C** and **Box 3** for primer design.

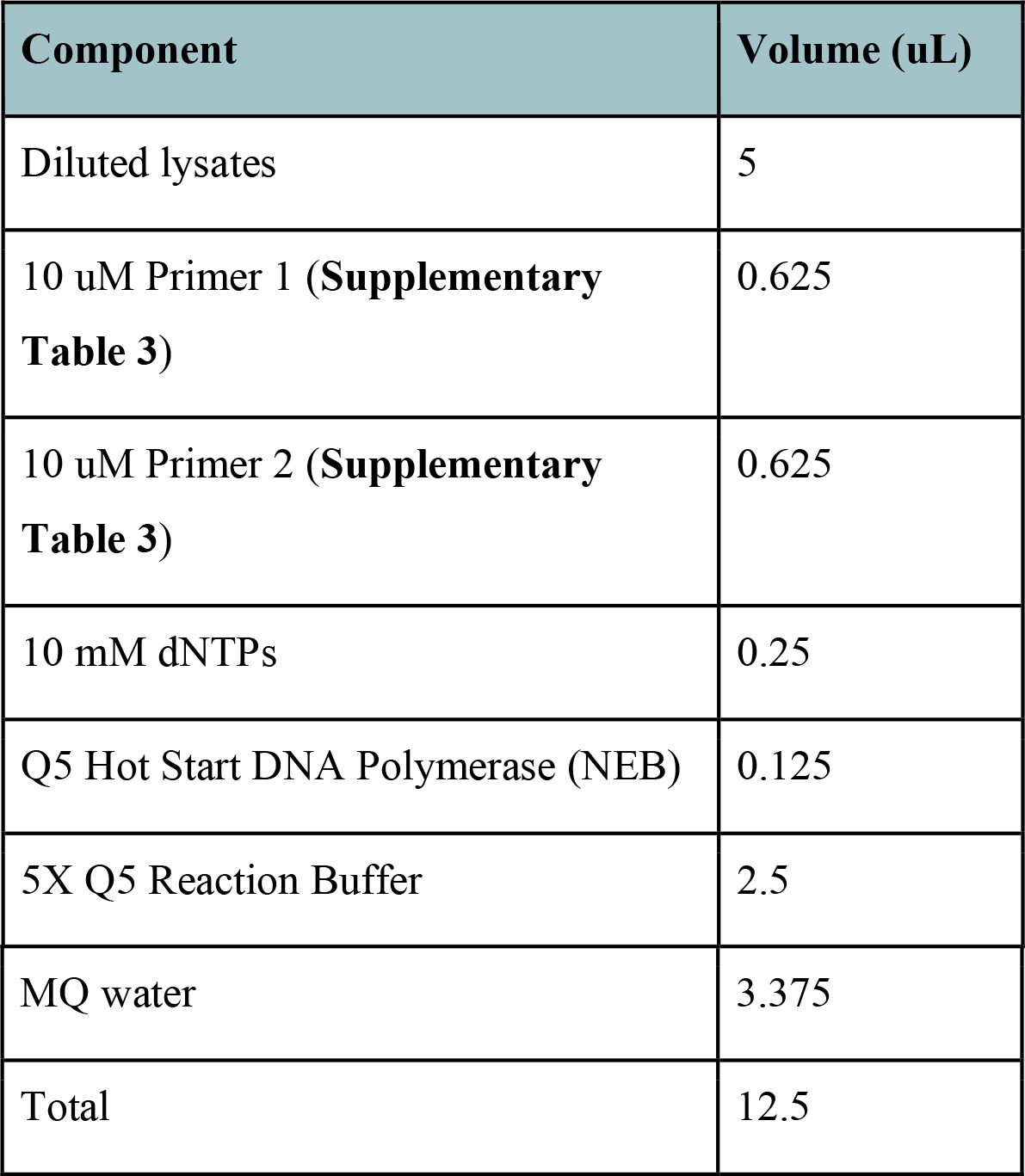
27) Perform PCR in a thermocycler with the following conditions:

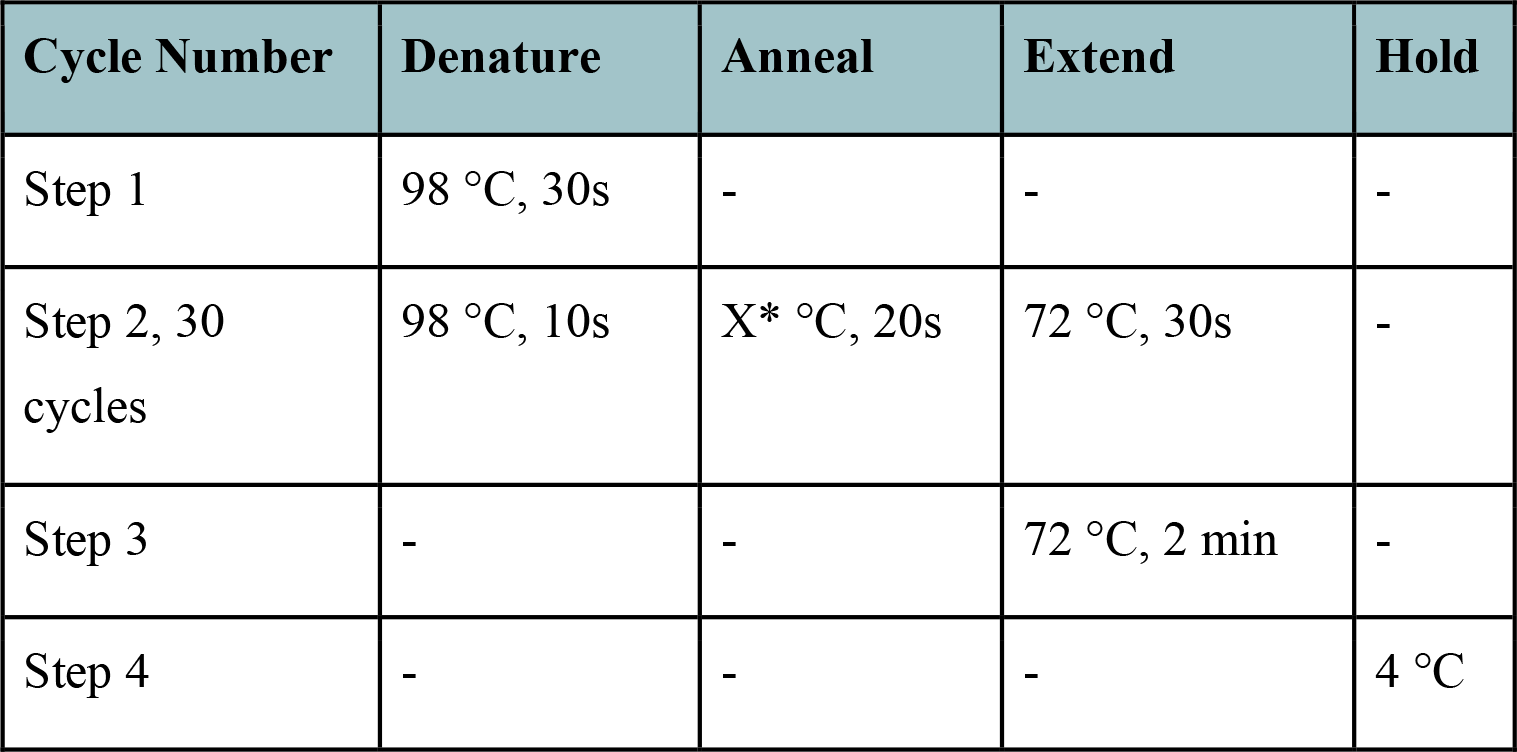 [CRITICAL] * Vary temperature of annealing step according to primer binding sites and the choice of polymerase. Annealing temperature estimates can be found online at tmcalculator.neb.com. Increase extension time for longer expected PCR products (for Q5 Polymerase, ∼30s of increased extension is recommended for each additional 1 kb of product).
28) Perform gel electrophoresis as described in Step 4. For PCR products smaller than 1 kb, a 1.5% agarose gel is recommended. Run electrophoresis so that the bands corresponding to inserted and uninserted products separate sufficiently (**Fig. 5D**).
29) Using a clean blade or razor, cut out the top inserted band for 2-3 clones with no uninserted product visible, and gel extract using the QIAquick or MinElute gel extraction kits (Qiagen).
30) Confirm insertion, including orientation and distance from target sequence, by Sanger sequencing of extracted bands. Either PCR primer (from step 26) can be used for sequencing. [TROUBLESHOOTING] Noisy Sanger sequencing chromatograms, see **Supplementary Table 4**.
31) Inoculate LB cultures from grown spots made in step 22 for downstream applications.

### (Optional) Construct plasmid curing - [Timing - 2-3 d]

32) Plasmid curing should be considered for clonal isolation before the start of the experiment. pSPIN plasmids on the pSC101* temperature-sensitive backbone can be cured efficiently by overnight growth at 37 °C (option A). Other plasmid backbones may also be cured by continued growth without antibiotic selection (option B); however, this method is not guaranteed to work across different plasmids and host cells, particularly for high-copy number plasmids (such as pUC19 pDonors).
  A. pSC101* plasmid curing.
    i. Into a 50 mL conical tube, inoculate 15 mL of LB media with no spectinomycin added using a cell spot from Step 23. Grow with shaking overnight at 37 °C. [CRITICAL] Antibiotics other than spectinomycin should still be added to the growth media if other plasmids or markers require active selection.
    ii. Dilute 2 uL of overnight culture into 500 uL of LB in a microcentrifuge tube. Plate 100 uL onto an agar plate.
    iii. Prepare 2-3 1:10 serial dilutions of diluted culture from Step 32Aii, each plated on a separate agar plate.
    iv. Incubate plates from Step 32Aii-iii at 37 °C overnight.
    v. After overnight incubation, choose a plate where single colonies are discernable. Pick 5-6 colonies, stamp each onto a new LB agar plate without spectinomycin and a new LB agar plate with 1X spectinomycin added. Number each stamp accordingly across the two plates.
    vi. Incubate plates overnight at 37 °C.
    vii. Stamped colonies growing only on the no-spectinomycin plate have had the pSC101* plasmid cured.
    viii. (Optional) PCR confirmation of plasmid loss. From the no-spectinomycin plate, pick colonies, lyse cells, and perform PCR and gel electrophoresis as in steps 23-28. Suggested primers are in **Supplementary Table 3**. Cells with cured plasmids should show no visible amplification.
  B. Curing of other plasmid backbones.
    i. Into a 50 mL conical tube, inoculate 15 mL of LB media using a cell spot from Step 23. [CRITICAL] LB media should not contain any antibiotics selecting for backbones that need to be cured from cells.
    ii. Grow with shaking overnight at 37 °C. A short growth (1-3 h) at 42 °C increases curing efficiency.
    iii. Plate cells and phenotypically characterize 20-30 colonies on selection plates as in step 32Aii-vi. Dilute 5 uL of the overnight culture from Step ii into 15 mL of fresh LB media, and incubate overnight at 37 °C. [CRITICAL] If curing more than one plasmid at once, colonies should be stamped on plates containing only one of the corresponding antibiotic types, in addition to plates without any antibiotics.
    iv. Isolate colonies growing only on the no-selection plate. If no such colony was observed, repeat Step iii using the new overnight culture from Step iii. For multiple plasmid curing, if a colony was found without one of the target curing plasmids, inoculate from that colony as in Step and repeat Steps 32Bi-iv.
    v. (Optional) PCR confirmation of plasmid loss. From the no-selection plate, pick colonies, lyse cells, and perform PCR and gel electrophoresis as in steps 23-28. Suggested primers are in **Supplementary Table 3**. Cells with cured plasmids should show no visible amplification.

### Anticipated results

After carrying out this protocol, the user should be able to generate a clonally integrated mutant that can be used for various applications (**Fig. 3**). We encourage systematic analyses in order to carefully assess the efficiency of both plasmid delivery and DNA integration, as well as optional analyses to interrogate genome-wide specificity by transposon-insertion sequencing (Tn-seq) (**Fig. 6**). For both heat-shock and electroporation techniques, transformation efficiency can be straightforwardly calculated by dividing the number of resulting colonies with vector backbone selection by the input amount of vector (20-100 ng recommended), taking the dilution factor used for plating into account (**Fig. 6A**). When conjugation is used as the delivery method and plasmids are stably maintained in recipient cells, efficiency can be calculated via selective plating on the plasmid vector backbone marker in comparison to plating without any selectable marker (**Fig. 6A**). The ratio of selection:no selection provides an estimate of how efficiently the plasmid was conjugated from the donor to the recipient strain.

**Fig 6.**
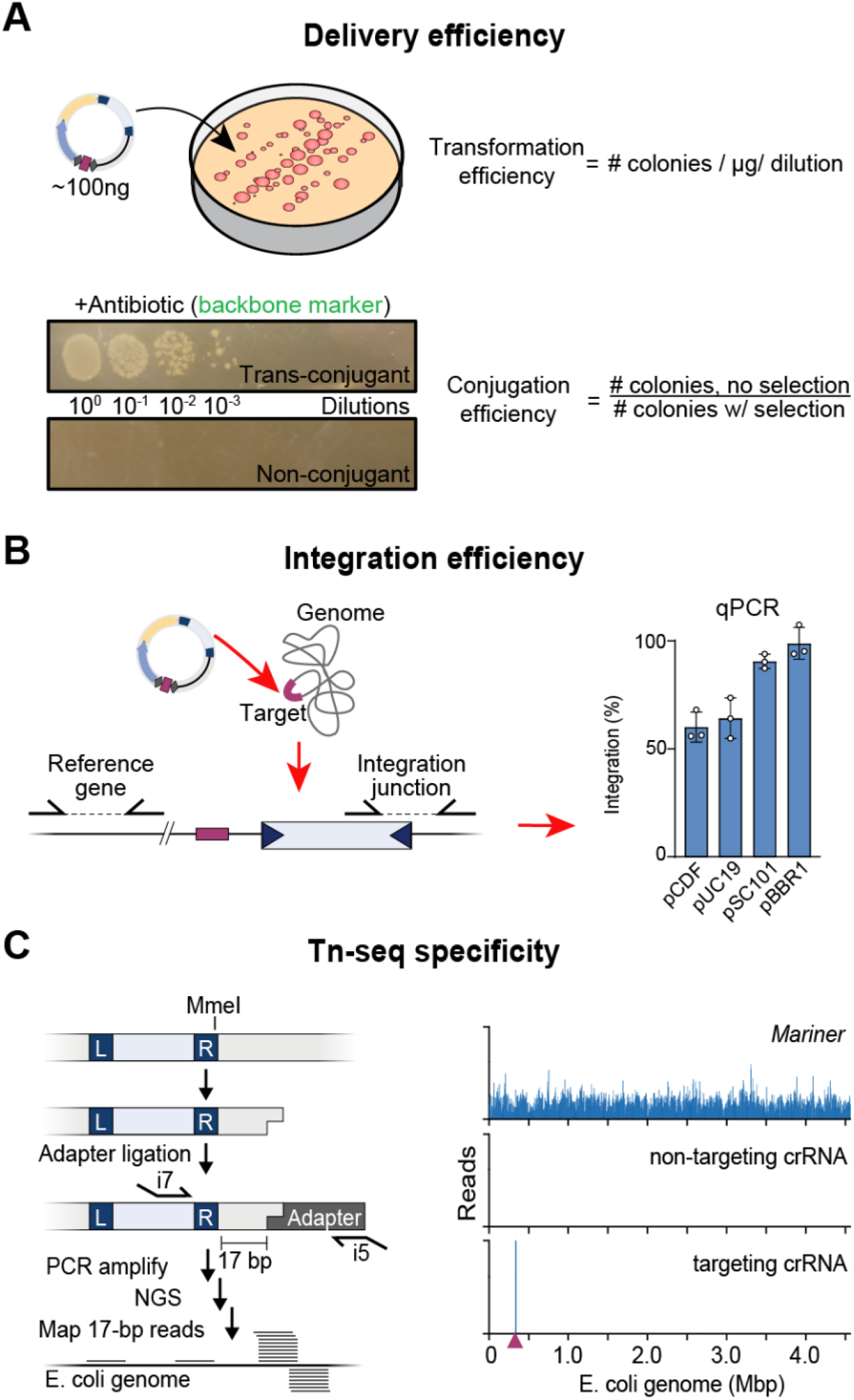
Anticipated results from a CAST editing experiment. (A) Examples and equations for calculating delivery efficiencies are shown for transformation (i.e. chemical or electroporation) and conjugation. (B) DNA integration efficiencies may be assessed and/or measured by PCR and/or qPCR. Representative data from experiments with pSPIN that were performed in *E. coli* are shown (right)^32^. (C) For users interested in profiling genome-wide specificity, Tn-seq may be applied to unbiasedly sequence all integration events within a sample of interest (left). Representative data for the use of *Mariner* or VchCAST in *E. coli* are shown at right, for both a non-targeting and targeting crRNA^33^.

As for the DNA integration reaction itself, we typically observe efficiencies of 50-99% for ∼1-kb payloads in *E. coli* and other bacteria (**Fig. 6B**), which can be calculated via qPCR of the integration junction and a reference gene locus (see **Fig. 5** for primer considerations). Excitingly, we have observed that these integration efficiencies can be further increased either by conducting transposition and growth at lower temperatures (30 C)^32^ or through the use of higher-activity homologous CAST systems^41^. Lastly, while beyond the scope of this protocol, it may be helpful to perform whole-genome sequencing on isolated clones to further confirm the genotype, as well as confirm the absence of off-target insertions, particularly when multiple genomic insertions of the transposon have been made during strain engineering. In previous studies, we performed Tn-seq using modified mini-Tn substrates with an MmeI digest site, in order to unbiasedly report on all genomic regions containing a mini-Tn insertion by high-throughput sequencing^31,33^ (**Fig. 6C**). This approach revealed that the VchCAST system typically integrates DNA payloads with >99% on-target specificity, as compared to the frequent non-specific insertions generated by both *Mariner* transposons and Type V-K CAST systems^32,33,41^. In addition, long-read sequencing on either the PacBio or Nanopore platforms can also provide confirmation of integration product purity (simple insertion vs cointegrate, **Box 1**) and specificity through individual reads spanning the entire transposon insertion^45,46^. However, since the high target-specificity of type I-F CAST systems are well documented with proper crRNA design^32,33,35,39^, many engineering applications may not require genome-wide analyses of transposition events.

In summary, CAST systems hold great value in genetically engineering both model and non-model bacteria, and hold potential for culture-independent editing of microbial communities. Microorganisms exist in dynamic complex communities that play fundamental roles that underpin every aspect of life, mediated by synergistic interactions between individual metabolic pathways and inter-microbial communication networks. Yet, our understanding of these interactions is sparse likely due to the limitations of the genetic tools used to determine gene functions, which are only applicable to pure cultures, where these complex interactions cannot occur. One key example of a microbiome that can benefit from CAST engineering is the human gastrointestinal tract, which comprises a complex and diverse microbial community whose composition and spatial architecture are increasingly being appreciated as critical drivers of human health and behavior^83^. The ability to directly manipulate the gut microbiome *in situ* is critical for mechanistic study of these microbial interactions in nature as well as develop novel therapeutics, yet the tools available for studying and modifying the complex gut microbial communities *in vivo* remain severely limited and insufficient. For example, high-throughput sequencing offers only observational information, germ-free mammalian systems poorly reflect natural host-microbiome interactions, and probiotics may offer novel genetic functions but suffer from limited temporal persistence in non-native environments. To address these key limitations, future work will require developing new platforms for precision microbiome engineering that combines programmable CASTs with broadly transmissible vectors for culture-independent microbial manipulation. These approaches should utilize CASTs to not only introduce edits, but also stably embed desired genetic payloads and allow for long-term persistence in microbiomes. These future thrusts in microbial engineering will define a new paradigm for genetic studies of microbiomes from diverse environments (i.e., gut, soil, ocean, extremes, etc.), enabling scientists to harness the power of microorganisms for the benefit of human advancement. Lastly, the application of CASTs to eukaryotic organisms, specifically mammals, holds enormous potential for genetic engineering of genes relevant to disease states.

## FUNDING

This research was supported by the National Institutes of Health (Grant numbers DP2HG011650 and R21AI168976 to S.H.S.), the Pew Biomedical Scholars Program (S.H.S.), the Alfred Sloan Foundation Research Fellowship (S.H.S.), and the Irma T. Hirschl Career Scientist Award (S.H.S.).

## Conflict of interest

Columbia University has filed a patent application related to this work for which P.L.V.., S.E.K., and S.H.S. are inventors. S.H.S. is a co-founder and scientific advisor to Dahlia Biosciences, a scientific advisor to CrisprBits and Prime Medicine, and an equity holder in Dahlia Biosciences and CrisprBits.

## Tables

**Supplementary Table 1. Description and sequence of plasmids.**

**Supplementary Table 2. All restriction enzyme digest sites on plasmid vectors.**

**Supplementary Table 3. All primer sequences and usages for experiments.**

**Supplementary Table 4. Troubleshooting guide for experiments.**

- Corresponds to all [TROUBLESHOOTING] in manuscript.

## Notes

### Competing Interest Statement

The authors have declared no competing interest.

